# A Hierarchical Model for eDNA Fate and Transport Dynamics Accommodating Low Concentration Samples

**DOI:** 10.1101/2024.03.27.586987

**Authors:** Ben C. Augustine, Patrick R. Hutchins, Devin N. Jones, Jacob R. Williams, Eric Leinonen, Adam J. Sepulveda

**Affiliations:** U.S. Geological Survey, Eastern Ecological Science Center, 12100 Beech Forest Road, Laurel, Maryland, 20708 USA; U.S. Geological Survey, Northern Rocky Mountain Science Center, 2327 University Way Suite 2, Bozeman, MT 59715, USA; Turner Institute of Ecoagriculture, Natural Resources Program, 901 Technology Blvd, Bozeman, Montana 59718

**Keywords:** environmental DNA, eDNA survival, eDNA transport, eDNA inhibitors, hierarchical modeling

## Abstract

Environmental DNA (eDNA) sampling is an increasingly important tool for answering ecological questions and informing aquatic species management; however, several factors currently limit the reliability of ecological inference from eDNA sampling. Two particular challenges are 1) determining species source location(s) and 2) accurately and precisely measuring low concentration eDNA samples in the presence of multiple sources of ecological and measurement variability. The recently introduced eDNA Integrating Transport and Hydrology (eDITH) model provides a framework for relating eDNA measurements to source locations in riverine networks, but little empirical work has been done to test and refine model assumptions or accommodate low concentration samples, that can be systematically undermeasured. To better understand eDNA fate and transport dynamics and our ability to reliably quantify low concentration samples, we developed a hierarchical model and used it to evaluate a fate and transport experiment. Our model addresses several low concentration challenges by modeling the number of copies in each PCR replicate as a latent variable with a count distribution and conditioning detection and quantification on replicate copy number. We provide evidence that the eDNA removal rate declined through time, estimating that over 80% of eDNA was removed over the first 10 meters, traversed in 41 seconds. After this initial period of rapid decay, eDNA decayed slowly with consistent detection through our farthest site 1km from the release location, traversed in 250 seconds. Our model further allowed us to detect extra-Poisson variation in the allocation of copies to replicates. We extended our hierarchical model to accommodate a continuous effect of inhibitors and used our model to provide evidence for the inhibitor hypothesis and explore the potential implications. While our model is not a panacea for all challenges faced when quantifying low-concentration eDNA samples, it provides a framework for a more complete accounting of uncertainty.

## Introduction

Environmental DNA (eDNA) approaches are increasingly being used to estimate ecological parameters like species distributions (Carraro et al., 2018), abundance (Rourke et al., 2022), and phenology (Searcy et al., 2022) due to their detection sensitivity, wide applicability across species, and cost efficiency, among other reasons (Jo and Yamanaka, 2022). Further, eDNA sampling can be a particularly useful tool for aquatic invasive species monitoring, potentially allowing for early detection and eradication (Larson et al., 2020; Morisette et al., 2021; Sepulveda et al., 2020). However, across many applications of eDNA monitoring in aquatic environments, the reliability of ecological inference can be reduced by 1) uncertainty in the source location of detected eDNA (Carraro et al., 2018; Jo and Yamanaka, 2022) and 2) difficulties measuring site and sample eDNA concentrations with minimal bias while accounting for all relevant sources of uncertainty (Ellison et al., 2006; Shelton et al., 2019; Espe et al., 2022). Both of these factors have the potential to reduce the effectiveness of eDNA sampling for management action, depending on their magnitude and how well they are addressed in experimental/monitoring design and statistical analysis.

A key requirement for many aquatic applications of eDNA sampling, particularly for invasive species monitoring, is to estimate source population locations because the location where eDNA is detected is not necessarily where it was produced due to hydrological transport (Carraro et al., 2018; Burian et al., 2021; Jo and Yamanaka, 2022). In stream or river networks, eDNA flows downstream from the source(s) and is detectable until the eDNA concentration attenuates below a detectable level due to physical degradation (Lance et al., 2017) and/or removal from the water column (Shogren et al., 2017). Therefore, an understanding of eDNA transport dynamics is required to either localize source populations (e.g., Carraro et al., 2018) or more broadly, estimate the plausible range of upstream distances a source population can be located from a detection to direct next steps, like more intensive sampling for confirmation (Sepulveda et al., 2023).

Environmental DNA fate and transport dynamics are the product of 1) ecological and biological factors determining how much eDNA is produced across source populations (e.g., abundance, biomass, eDNA production rate), 2) hydrological factors (e.g., discharge, particle settling and resuspension, river network connectivity), and 3) their interaction (e.g., eDNA degradation, removal from water column to substrate) (Carraro et al., 2018; Curtis et al., 2021; Troth et al., 2021; Shogren et al., 2017). A recent modeling framework that includes these general features is the eDNA Integrating Transport and Hydrology (eDITH) model (Carraro et al., 2018, 2021), which has been used to predict the locations of source populations and their eDNA production rates (Carraro et al., 2021) that can correlate with abundance (Jo and Yamanaka, 2022). The eDITH model is necessarily simplistic, given the number of factors that need to be accounted for with the typical level of eDNA sampling and the need to apply the model to large river networks which can be computationally intensive. Further, the eDITH model likelihood can be multimodal (described as “equifinality” by Carraro et al., 2021), with the eDNA removal rate parameter(s), the product of both physical decay and removal, being poorly estimated (Carraro et al., 2021). Therefore, prior information, particularly about the eDNA removal rate parameter(s), can improve parameter estimation and thus ecological inference (Carraro et al., 2021, 2023).

Release experiments (e.g., Jane et al., 2015; Laporte et al., 2020) have been important for improving our understanding of eDNA transport and removal dynamics, and they offer a means of providing parameter estimates that can be used as prior information in future modeling when the source locations are not known in advance (e.g., Carraro et al., 2018). Further, release experiments allow us to test model assumptions (Bylemans et al., 2018; Yates et al., 2021), better evaluate the level of realism necessary for reliable inference, and assess the data demands for a given level of realism. For example, eDITH model applications to date have assumed that the eDNA removal rate is constant with respect to time, which could lead to biased estimates of source locations or detection distance from a source if violated. In fact, the eDNA removal rate has been shown to decline with time in some experiments (Bylemans et al., 2018; Yao et al., 2022; Snyder et al., 2023). Bylemans et al. (2018) hypothesized this decline may be due to variable removal rates across eDNA fragments, such as between fragments in cells and free-floating DNA, and Snyder et al. (2023) provided experimental evidence that removal rate varies by particle size. To our knowledge, these hypotheses have not been compared to empirical data within the eDITH framework, which is possible in release experiments if the relevant hydrological variables are measured accurately.

Another key requirement of many aquatic applications of eDNA sampling is the accurate and precise quantification of eDNA concentration. While many ecological questions can be addressed with eDNA detection data alone (e.g., Hunter et al., 2015), relating eDNA measurements to abundance or modeling eDNA production, transport, and removal as a function of hydrology require quantitative eDNA measurements. These measurements can be obtained by quantitative or digital PCR (Yates et al., 2019), where measurements are made across multiple PCR replicates (hereafter ‘replicates’) of the same sample. These replicate-level measurements are typically then summarized and treated as data, usually using the mean concentration across replicates as the sample concentration or averaging replicates across samples to produce site concentrations (e.g., Yates et al., 2021). However, replicate measurements vary due to multiple factors including variable concentrations across samples (Chambert et al., 2018; Shelton et al., 2019), variability in the allocation of eDNA copies to replicates (Rossmanith and Wagner, 2011; Dorazio and Hunter, 2015; Tellinghuisen, 2020), and measurement error in replicate copy number/concentration (Shelton et al., 2019; Espe et al., 2022). The common approach of averaging replicate concentrations does not partition multiple sources of variance and pools the measurement error with the ecological variance, potentially eroding ecological inference if the measurement error is non-negligible in magnitude relative to ecological variation.

Low concentration samples present several unique challenges for accurate eDNA quantification. First, eDNA copy number or concentration estimates are typically modeled assuming approximate normality (usually on the log scale, see Carraro et al., 2018; Espe et al., 2022), which is an accurate approximation at high concentrations where copy numbers in replicates are large, but less so as concentrations decline and there are fewer copies per replicate (Ellison et al., 2006; Dorazio and Hunter, 2015). Further, copy number measurements are often censored from below by a fixed number of quantitative (q)PCR cycles or a limit of quantification, which requires lower truncation of the observation model distribution (Espe et al., 2022). Low concentration samples also present challenges for interpreting nondetections—as sample concentrations decline, both true negative (Poisson sampling zeros) and false negative (failed detections) replicates become more likely, which cannot be deterministically separated given the observed data (Ellison et al., 2006). In this situation, excluding zeros from sample means will introduce positive bias, while including them may also introduce bias, with the direction depending on the relative proportion of true and false negatives. Finally, sample concentration can be underestimated due to interference by eDNA inhibitory compounds in water samples (i.e., inhibition), which can be difficult to reliably detect (Kontanis and Reed, 2006; Lance and Guan, 2020), and these inhibitors are more likely to affect lower concentration samples (McKee et al., 2015; Hunter et al., 2019). Such underestimation can bias ecological inference–for example, the eDNA removal rate in a release experiment would be overestimated or the source location prediction from an eDITH model would be too far away. These challenges for quantifying low concentration samples are of particular concern when the goal is the early detection of invasive species, which are easiest to eradicate when population sizes are small, yet those populations produce less eDNA, yielding more samples where quantitative measurements are unreliable.

Hierarchical models are useful for partitioning multiple sources of variation and reducing bias by allowing the source(s) of bias to be modeled more mechanistically (Royle and Dorazio, 2008). These models are increasingly being used in eDNA analyses–for example, multiscale occupancy models (Nichols et al., 2008; Dorazio and Erickson, 2018; Stratton et al., 2020) have been used to propagate uncertainty in replicate-level occupancy states to site occupancy estimates and covariate relationships, and occupancy models have been extended to account for false positives (e.g., Smith and Goldberg, 2020; Guillera-Arroita et al., 2017). Hierarchical models have been used for quantitative eDNA data to partition measurement and multiple levels of environmental variation and relate both detection and quantitative measurements to site concentration (Shelton et al., 2019; Espe et al., 2022). However, to deal with the challenges of low concentration eDNA measurement, a hierarchical model representing the replicate copy numbers as latent discrete random variables may provide improved inference. In such a model, the copy numbers can be assigned a more appropriate count distribution, and the uncertainty in whether observed zeros are true or false negatives can be propagated to the ecological parameters of interest. Further, representing the copy numbers directly in the model may improve our ability to assess lack of fit at the replicate level and investigate hypotheses about inhibitors affecting the replicate-level measurements.

To better understand eDNA transport/removal, environmental variation, measurement error, and our ability to jointly estimate the parameters of these processes, we conducted an eDNA release experiment and developed a hierarchical model to apply to our experimental data that accommodates all three sources of nondetections described above. Of particular concern was how well the eDITH model parameters are estimated and whether the eDNA removal rate is constant or declining through time. Because we observed that our original model did not adequately fit our experimental data, particularly for lower concentration samples, we hypothesized that the source of poor fit could be eDNA inhibitory compounds and expanded the model to accommodate the impact of eDNA inhibitors on detection and copy number quantification. We used this model to illustrate how inhibitors can bias inference of multiple parameters and demonstrate how they can, in principle, be accommodated through hierarchical modeling for more reliable ecological inference.

## Field Methods

### DNA release

We conducted this experiment in the upper section of Green Hollow Creek, a 1st-order tributary of the Gallatin River located on the Flying D Ranch in southwest Montana (USA), 2023 August 21-24. The upper section of this stream is approximately 1 m wide, 3-12 cm deep, a gradient of 50 m per km, had a mean discharge 0.02 cms during our study and flows into a small reservoir containing a conservation brood stock of Arctic grayling (*Thymallus arcticus*), the target species for this study. An artificial barrier restricts upstream movement of Arctic grayling from the reservoir into the Green Hollow Creek study reach– this species is absent in the watershed upstream of the barrier. We introduced water collected from the reservoir’s offshore zone to the injection point (0 m) at the top of the 1000 m study reach. This experimental water was transported from the reservoir in *∼*200 L carboys and then transferred into bleached and rinsed coolers connected in series and placed in a shaded area adjacent to the injection point. The water was refreshed every 12 hr. Beginning at time 0 hr, water was dripped into the stream at a rate of 220 mL/min for 49 hr using an electric-operated pump connected to the cooler series and 2.5-cm PVC that spanned the width of the stream. Eight sprinkler emitters were attached to the PVC at 0.1 m intervals so that experimental water was dripped across the stream’s width to facilitate mixing. At time 32 hr, eight Arctic grayling that were recreationally-angled and legally harvested were added to the cooler series. Experimental water was dripped into the stream until eDNA sampling concluded.

### eDNA Sampling

At time 48 hr, we used distinct 19-L buckets to collect eDNA samples at -0.5, 0.5, 2.5, 5, 10, 20, 40, 80, 125, 200, 300, 400, 500, 1000 m from the injection point. Sampling was completed at all sites within 5 min. Buckets were bleached (50% commercial bleach, Goldberg et al., 2016), rinsed with stream water upstream of the injection point, and then repeatedly rinsed with stream water at the sampling point prior to collecting a sample. To collect samples, we filled the bucket with stream water from the cross-section midpoint, swirled the bucket, and then poured this water into each of six, 1-L sterile whirlpak bags. Our intention with the buckets was to collect uniform samples at and across sampling points in as short a time-increment as possible to minimize any temporal variability associated with inconsistent eDNA production and removal rates. Bags were stored at ambient temperatures in the shade and filtered on site through 47-mm diameter, 1.2 µm mixed cellulose ester filters (Millipore) within 90 min of collection. Filters were placed immediately into 200 µL lysis buffer, which contained 20 µL of proteinase K and 180 µL of Qiagen Buffer AE, and returned to USGS Northern Rocky Mountain Research Station (NOROCK) for extraction and analysis.

We collected two field negative controls immediately before time 48 hr, which consisted of 250-ml of deionized water poured into sterilized whirlpak bags. We collected field positive controls by sampling 1 L of water directly from the reservoir at time -24 and 0 hr, and by sampling 1 L of water from the coolers at time 0, 24, and 48 hr. Controls were handled similarly to experimental samples. Negative controls were filtered immediately prior to experimental samples, whereas positive controls were filtered at USGS NOROCK.

### Hydrological Covariates

Discharge was measured at all eDNA sample collection sites with the following exceptions due to stream habitat complexity (e.g., undercut banks, large wood and cobbles) that prevented accurate measurement: we used discharge measured at 5 m for the first three sites at 0.5, 2.5 and 5 m; we used discharge measured at 15 m for sites at 10 and 20 m; and we used discharge measured at 35 m for the site at 40 m. Discharge measurements were estimated using the velocity-area method, which involves dividing each site cross-section into multiple subsections and measuring the width, depth, and flow velocity of each subsection with a hand-held current meter. Discharge values for each subsection were summed to arrive at the cross-section discharge. Discharge was sampled at time 24 hr and time 49 hr. We also deployed barometric pressure transducers (Onset HOBO water level logger) set to 1-hr intervals in the air and water at 125 m to evaluate whether the water surface elevation (as a proxy for discharge) was stable throughout our experiment. The transducer in the water also collected water temperature at hourly increments. We deployed additional water temperature loggers (Onset HOBO pendant) at -0.5 and 1000 m to describe water temperature at hourly increments.

## Lab Methods

### Assay Development

Assays designed for eDNA studies using qPCR generally target short amplicons that are between 50-150 base pairs (bp) in length (Goldberg et al., 2016; Rees et al., 2014) because PCR efficiencies are higher for shorter amplicons (Bustin and Hugget, 2017) and because shorter DNA fragments tend to be more available for detection in aquatic environments than longer fragments (Bylemans et al., 2018). To explore differences in transport dynamics of short and long DNA fragments, two probe-based qPCR assays were developed for use in this study. Both assays target the *T. arcticus* cytochrome oxidase subunit 1 gene (cox1) and share the same reverse primer. The first unique set of forward primer and positive-sense strand probe produces a 128 bp amplicon while the second set produces a 468 bp amplicon. *In silico* validation was performed by aligning all available *T. arcticus* cox1 gene from NCBI and BOLD databases and identifying conserved regions of DNA greater than 15 base pairs in length. We then used NCBI’s nucleotide BLAST to remove areas with high similarity to any off-target sequences. The primer sequences that were selected with the appropriate thermal properties minimized similarity with sequences outside of the Thymallus genus. The only teleost fish (outside of the Thymallus genus) with fewer than three base pair mismatches with at least one primer was *Liopropoma olneyi*, a Caribbean reef fish. NCBI Primer-BLAST was used to confirm this result in the nucleotide (nr) database of teleost fishes using default stringency and specificity settings. It is important to note that, while the assays designed here can reliably detect *T. arcticus*, there is also significant sequence similarity with other members of the Thymallus genus. Therefore these assays should not be considered species specific when used in water bodies that may contain other members of Thymallus.

Each assay’s annealing temperature was optimized using temperature-gradient qPCR (from 58-67 °C), wherein the optimal annealing temperature was that which minimized the Cq when amplifying replicate synthetic DNA standards of 5000 gene copies. The optimal primer ratios were determined similar to Wilcox et al. (2015). The limit of detection (LOD) and limit of quantification (LOQ) were determined in both an ideal sample matrix (i.e., molecular grade water) and in a relevant sample matrix (extracted water from Green Hollow Creek) using the eLowQuant method of Lesperance et al. (2021).

The assays were validated *in vitro* with DNA extracted from *T. arcticus* from Green Hollow Creek and from tissues of regional origin of *Salvelinus fontinalis*, which do occur in our study reach, and other salmonids found in nearby waters including *Oncorhynchus mykiss, O. clarkii bouvieri, O. clarkii lewisi, Salmo trutta*, and *Prosopium williamsoni*. Tissue validation was performed with 10 replicates each. *In situ* validation was performed on eDNA water samples collected from the Green Hollow brood stock reservoir (known positive source) and from Green Hollow Creek water collected above the barrier prior to the start of the experiment (known negative source).

### DNA Extraction

All eDNA and tissue samples were extracted with Qiagen DNeasy Blood and Tissue Kits (cat. #69504) with Qiagen DNeasy Lyse and Spin Baskets (cat. #19598) with minimal alterations of the manufacturer’s protocol. DNA was eluted in 400 µL of Qiagen Buffer AE. All DNA extraction batches included an extraction negative control using only extraction kit reagents. These extraction blanks were otherwise handled and analyzed identically to samples.

### Quantitative PCR

Assays were run on a BioRad CFX 96-Touch or BioRad Opus thermal cycler (Hurcules, CA). Reactions took place in BioRad Hard-Shell® optical 96-well plates (cat. #HSP9601) sealed with BioRad Microseal® optical adhesive film (cat. #MSC1001). The thermal cycle was: 95 °C for 15 minutes then 50 cycles of 94 °C for 15 seconds and 60 °C for 60 seconds. Reactions of 20 µL included 10 µL of Qiagen Quantitect Master Mix, 3 µL sterile water, 0.5 µM of forward and 0.4 µM of reverse primer, 0.25 µM FAM-labeled hydrolysis probe, and 4 µL of DNA extract. The two *T. arcticus* assays were not multiplexed together, but were run in separate reactions multiplexed with an internal positive control (IPC, see below). Every 96-well plate run as part of this study had the same duplicated (two replicates) controls: six point 1:10 dilution *T. arcticus* synthetic standard curve ranging from 4e0 to 4e5 target DNA copies, no-template control (NTC), and internally blocked negative control (IBC). FAM and VIC fluorescence values were ROX-normalized prior to baseline correction and amplification detection (Patrone et al., 2020) using R v4.0.3 statistical software (R Core Team, 2021). Successful amplification of DNA was defined as any curve for which a feasible solution with less than 10% transformation error was found for the non-linear optimization transformation presented in Patrone et al. (2020).

### Inhibition Assessment

Inhibition was measured through the use of an exogenous IPC in every qPCR reaction, including controls (TaqMan™ Exogenous Internal Positive Control Reagents, cat. #4308321). We added 2 µL of VIC-labeled 10x Exo IPC Mix and 0.4 µL 50x Exo IPC DNA per qPCR reaction. Standard dilutions of IPC DNA were not used, so inhibition was assess using quantification cycle (Cq) values determined from the methods described and referenced above. We used ΔCq to describe inhibition of each qPCR reaction on a continuous scale:

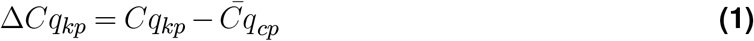

where Δ*Cq*_*kp*_ is the difference in IPC *Cq*_*kp*_ for qPCR reaction *k* on plate *p* and the mean Cq of a set of *c* uninhibited control samples on plate 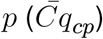. Said another way, Δ*Cq*_*kp*_ is the difference between a sample IPC Cq and the mean IPC Cq of a set of uninhibited samples on the same 96-well plate. Uninhibited control samples for each plate included no-template controls (NTCs) and synthetic standards with copy numbers of the target DNA strand <1000. Positive values of Δ*Cq*_*kp*_ reflect a delay in Cq, an indication of possible inhibiting effects.

## Modeling Methods

### Data Description

Our eDNA detection and quantification data are structured by site, sample, and PCR replicate. Sites are numbered *i* = 1,…, *I* = 13, each with a distance from the release site, *d*_*i*_, in sequential order starting at 0.5m. At each site, samples are numbered *j* = 1,…, *J* = 6, and for each sample, replicates are numbered *k* = 1 …, *K* = 5. We define 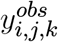 to be the measured copy number for site *i*, sample *j*, and replicate *k*, with copy number measurements converted from the observed Cq values. Elements of ***y***^*obs*^ corresponding to failed detections are set to 0 as are any copy number observations less than a lower bound for quantification, *η*^*q*^. This quantification lower bound can be set to the number of PCR cycles used to account for data censoring due to a limited cycle number (Espe et al., 2022), or it can be set higher to exclude the lowest copy number measurements if they do not meet model assumptions (e.g., biased measurement at low concentrations, discussed below).

Next, we define 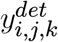 to be the detection data taking value 1 if the copy number measurement is greater than *η*^*d*^ and 0 otherwise, where *η*^*d*^ is a lower bound of detection that is less than or equal to the lower bound of quantification. For our analyses, we set *η*^*q*^ to the copy number corresponding to 50 cycles and we set *η*^*d*^ = 0 because we expect every sample to contain target eDNA. If false positives are a concern, both *η*^*d*^ and *η*^*q*^ can be set higher. Note, these definitions of detection and quantification lower bounds differ from standard definitions of “limit of detection” and “limit of quantification” (e.g., Klymus et al., 2020). Finally, for the hydrological data, we define *Q*_*i*_ to be the discharge rate at site *i* and *v*_*i*_ to be the average stream velocity between the release location and site *i*. Due to concerns about the measurement precision of stream velocity at each site relative to typical variation over a 1 km stream segment, we set the average stream velocity between the release location and each site to the mean of measured velocities across sites: 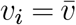.

### Ecological Process Model

#### eDNA Production, Transport, and Removal Process

We assume site eDNA concentrations are the product of sub-processes for eDNA production, transport, and removal for which we adapt the eDNA Integrating Transport and Hydrology model (eDITH; Carraro et al., 2018). The eDITH model is:

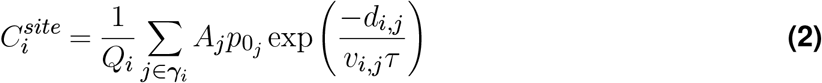

where 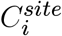 is the concentration at site *i* (*N/m*^3^), *Q*_*i*_ is the water discharge rate at site *i* (*m*^3^*/s*), ***γ***_*i*_ is the set of sites upstream of site *i, A*_*j*_ is the habitat area of site *j* (*m*^2^), *p*0_*j*_ is the eDNA production rate of site *j* (*N/m*^2^*s*), *d*_*i,j*_ is the distance between site *i* and *j* (*m*), *v*_*i,j*_ is the mean velocity between site *i* and *j* (*m/s*), and *τ* is the inverse eDNA removal rate (*s*). In our experiment, eDNA is added at a point location of zero area, so we remove area and define *p*_0_ to be the eDNA production rate without respect to area with units *N/s*. Further, eDNA is only “produced” at the release location, removing the need to sum the eDNA contribution of multiple upstream sites. These modifications simplify the eDITH model to:

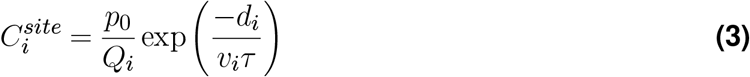

where *p*_0_ is the eDNA production rate at the release location, *d*_*i*_ is the distance between site *i* and the release location, and *v*_*i*_ is the mean velocity between site *i* and the release location, which we assume to be constant across sites.

We consider two versions of the eDITH model above with respect to how eDNA is removed from the water column as a function of time. Note that the expected travel time from the release location to site *i* is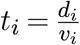, and the exponential term in the eDITH model is an exponential survival function in continuous time:

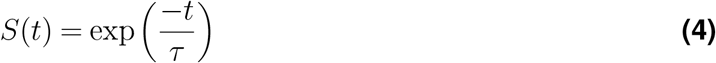

Note that the removal rate is *F* (*t*) = 1 *− S*(*t*), and we will use this term for better consistency with the eDNA literature. Under an exponential model, DNA is removed at the same rate through time. A Weibull model (David and Mitchel, 2012; Bylemans et al., 2018) considers that the eDNA removal rate can increase or decline through time, but includes an extra parameter, which we found led the eDITH model parameters to be weakly identified with our data. A power law relationship (Shogren et al., 2017; Levi et al., 2019) allows the eDNA removal rate to decline through time without an extra parameter, so we consider both an exponential and power law removal model. The power law survival function is:

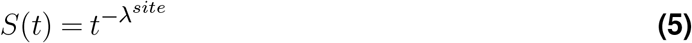

The resulting eDITH model with power law eDNA removal in continuous time is then:

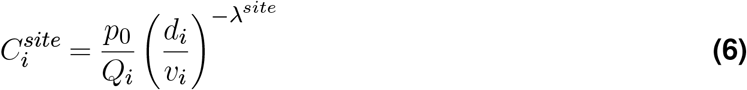

for sites 2,…,13. For both eDITH removal models, we add a thinning parameter at the first site to account for the plume effect seen in previous studies (e.g., Laporte et al., 2020; Wood et al., 2021), for example, in the power law version:

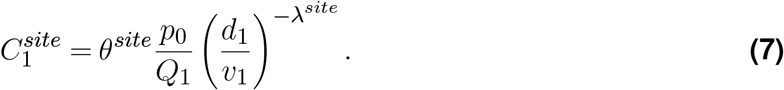

where 0 < *θ*^*site*^ < 1. As a consequence, copy number measurements from site 1 do not contribute to the estimation of the eDITH model parameters beyond ensuring that the concentration at site 2 is greater than or equal to that at site 1.

#### Sampling Process

The sampling process describes the variation in sample concentrations collected at a site as a function of the site concentration. This distribution is typically right-skewed due to eDNA clumping in the water column and stochastic collection of more rare, larger “aggregate” particles (Furlan et al., 2016; Yates et al., 2023); therefore, we use a lognormal distribution (Carraro et al., 2018; Espe et al., 2022) to describe this variation. Conditional on the eDNA concentration at site *i*, we assume that the concentration in each collected sample, 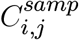, varies following:

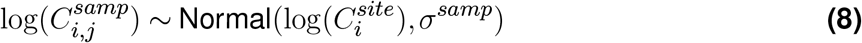

where *σ*^*samp*^ is the sampling process standard deviation on the log scale, which could be a function of site covariates such as the distance from the release location.

#### Replication Process

The replication process describes the variability in the number of copies allocated to each replicate given the sample concentration. We model the eDNA copy number in each replicate, *N*_*i,j,k*_, as a count random variable with an expected number of copies being a function of the sample concentration (Dorazio and Hunter, 2015; Furlan et al., 2016). More specifically,

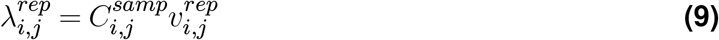

where 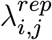 is the expected number of copies in a replicate from site *i* and sample *j* and 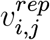 is the sample volume associated with 1 replicate from site *i* and sample *j*. If this volume varies across replicates of the same sample, a replicate dimension can be added to ***λ*** and ***v***^*rep*^. Next, we assume that the eDNA copies are homogeneously distributed throughout the eDNA extract and deposited into each replicate following a Poisson distribution, which has theoretical support for both digital PCR (dPCR) and qPCR (Dube et al., 2008; Rossmanith and Wagner, 2011; Tellinghuisen, 2020; Lesperance et al., 2021):

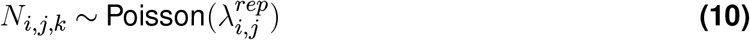

#### Observation Model

The observation model describes both the detection and quantification processes. For the detection process, we assume replicate-level detection probability, 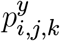, is a function of the number of copies in a replicate:

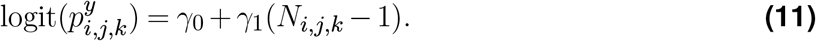

By subtracting 1 from the number of copies in each replicate, *γ*_0_ corresponds to the detection probability of 1 copy. Then, we assume

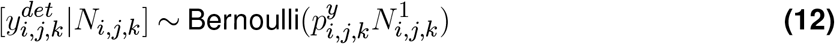

where 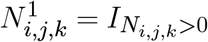 is an indicator variable that zeroes out the detection probability when 0 copies are allocated to a replicate on the logit scale. An alternative specification (Furlan et al., 2016) is:

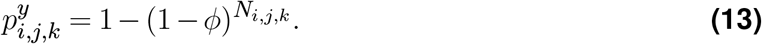

where *ϕ* is the 1 copy amplification probability and detection always occurs if one or more copies, acting independently, amplify. However, if the probability of detection depends on the number of copies that amplify, which could occur due to difficulty distinguishing positive detections from background signals (Forootan et al., 2017; Hunter et al., 2017), this model may not be appropriate.

Finally, the quantification process, conditioned on detection and the number of copies in a replicate, is

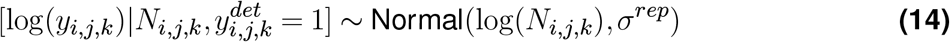

where *y*_*i,j,k*_ is the measured replicate copy number and *σ*^*rep*^ is the replicate-level copy number measurement error on the log scale, which could be a function of site, sample, or replicate covariates. Because 1) we condition the quantification process on positive detections and 2) copy number measurements may be censored from below by *η*_*q*_ (Espe et al., 2022), the observed replicate-level quantitative data, 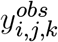, are:

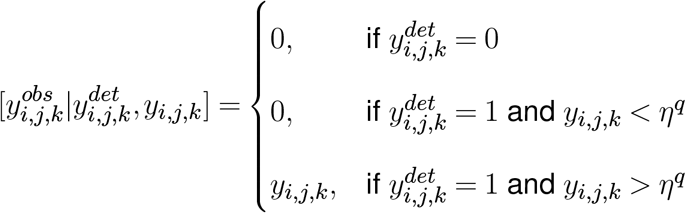

This model is depicted in Figure 1. An important assumption in this quantification process model is that the log copy number measurements are unbiased, i.e., *E*[log(*y*_*i,j,k*_)] = log(*N*_*i,j,k*_), which can be violated, for example, by a concentration plateau effect (Hunter et al., 2017) or by DNA inhibitors (McKee et al., 2015; Hunter et al., 2019; Sepulveda et al., 2020). We consider the latter in the next section.

**Figure 1.**
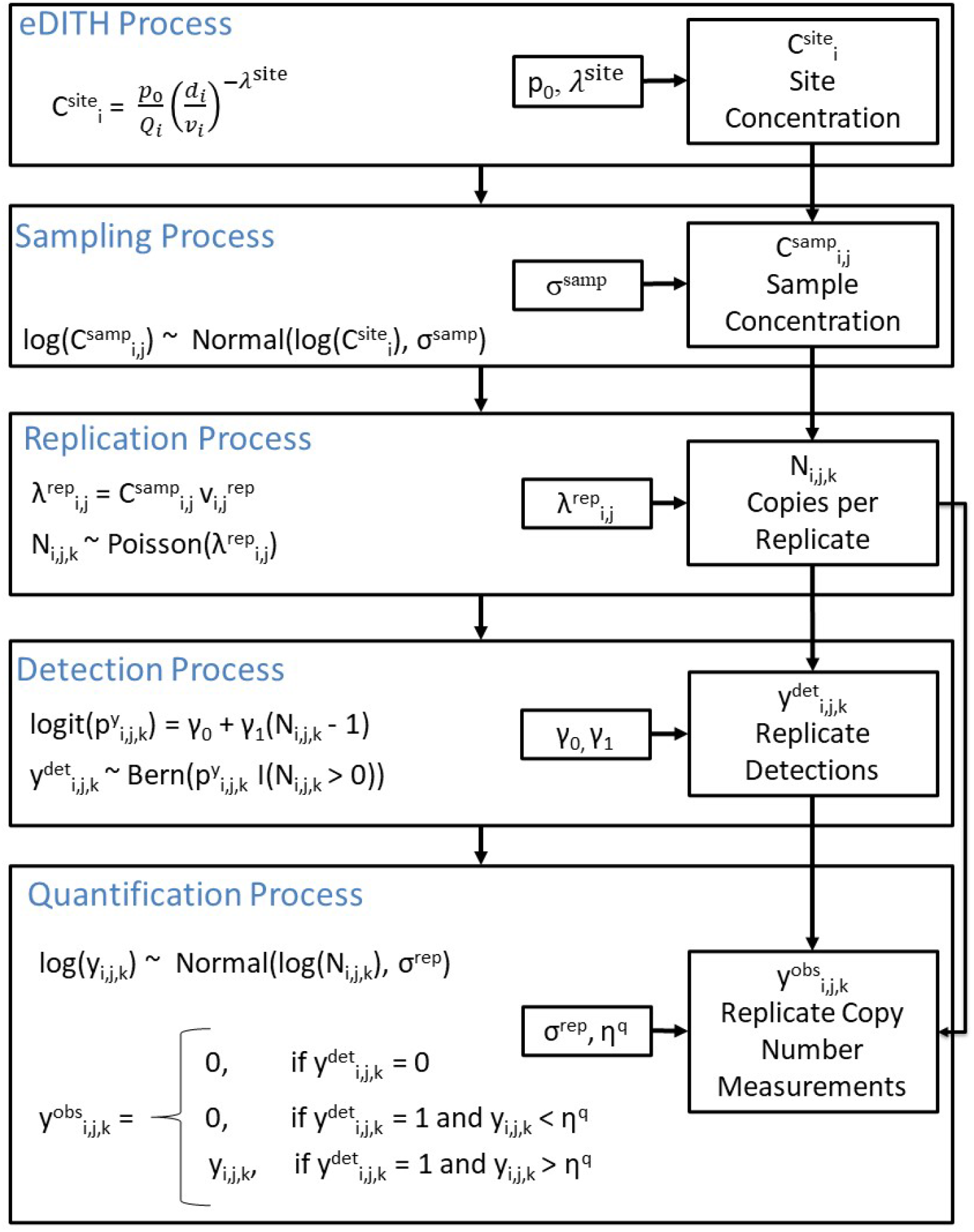
Model diagram without inhibitor processes. The eDITH process describes how concentration is distributed across sites, the sampling process describes how concentration is distributed across samples at a site, the replication process determines how many copies are allocated to each replicate for each sample, the detection process describes how copies in each replicate are detected, and the quantification process describes how detected copies are measured.

### Extended Process Model - Copy Inhibition

Here, we consider that the PCR reaction in each replicate may be partially or completely inhibited, which can decrease both the detection probability and the number of copies measured given detection. Compounds found in the environment can inhibit PCR reactions via multiple mechanisms, including binding to the target DNA and binding to or otherwise interfering with reaction products (Opel et al., 2010; McKee et al., 2015). Inhibition is typically identified by comparing the Cq values of IPCs to uninhibited control samples (Volkmann et al., 2007; Jane et al., 2015, see Lab Methods - Inhibition Assessment) and using a deterministic decision rule to exclude samples likely to be inhibited (e.g., samples with a ΔCq ≥ 3 Goldberg et al., 2016), though others have treated ΔCq as continuous measures of inhibition (Volkmann et al., 2007; Lance and Guan, 2020). We take the latter approach, modeling the probability of inhibition as a function of replicate copy number, replicate ΔCq, and a sample-level random effect. For each replicate, the probability of inhibition is 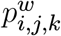, where

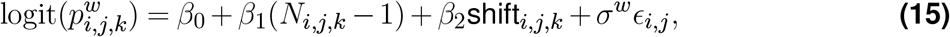

shift_*i,j,k*_ is the standardized ΔCq covariate, and *ϵ*_*i,j*_ *∼* Normal(0, 1) (non-centered sample-level random effects; Papaspiliopoulos et al., 2007). The sample random effects account for sample-level heterogeneity in replication inhibition probability, due, at least in part, to sample-level variability in inhibitor concentration. Next, we assume replicate inhibition states are Bernoulli random variables

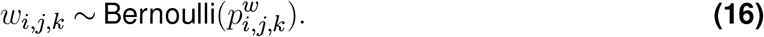

with *w*_*i,j,k*_ taking value 1 when inhibited and 0 otherwise. If a replicate is inhibited, we assume the copies available to be detected and measured, 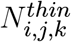, result from a thinning process of the true copy number:

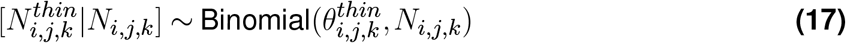

where 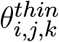 is the thinning rate, which could be a function of covariates or random effects. For simplicity, we assume 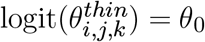. Then, to determine how many copies are available for detection and measurement as a function of the replicate inhibition states, we specify

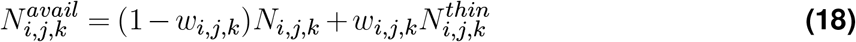

which evaluates to 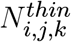 if a replicate is inhibited and *N*_*i,j,k*_ otherwise. Finally, we replace *N*_*i,j,k*_ with 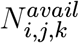 in the observation model so that detection and measurement are both conditioned on the inhibition states:

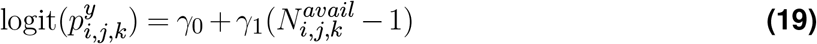

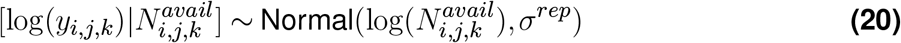

Then, the detection model is the same as equation 12, except we redefine 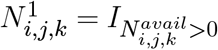, a function of the available copy number instead of the true copy number.

The number of copies available to be detected can be interpreted as the number of copies that inhibitors did not bind to, the number of copies measureable given the magnitude of binding of inhibitors to polymerase in a replicate, or a combination of both factors. More generally, the main feature of the inhibitor model is a large reduction in the detection probability and measured copy number for inhibited samples, which happens regardless of the precise mechanism (Opel et al., 2010; McKee et al., 2015). This extended model is depicted in Figure 2. With this model structure, if a replicate is inhibited, its detection probability and expected log copy number measurement given detection both decrease, with detection probability decreasing to 0 if no copies are available to be detected. Note that the inhibition states, *w*_*i,j,k*_, are not directly observed–we can only observe the effects of inhibition on the copy number detections and measurements. However; the specification of our inhibitor model as part of a larger hierarchical model (Royle and Dorazio, 2008) allows the inhibition states to be estimated jointly along with the other model variables and latent states.

**Figure 2.**
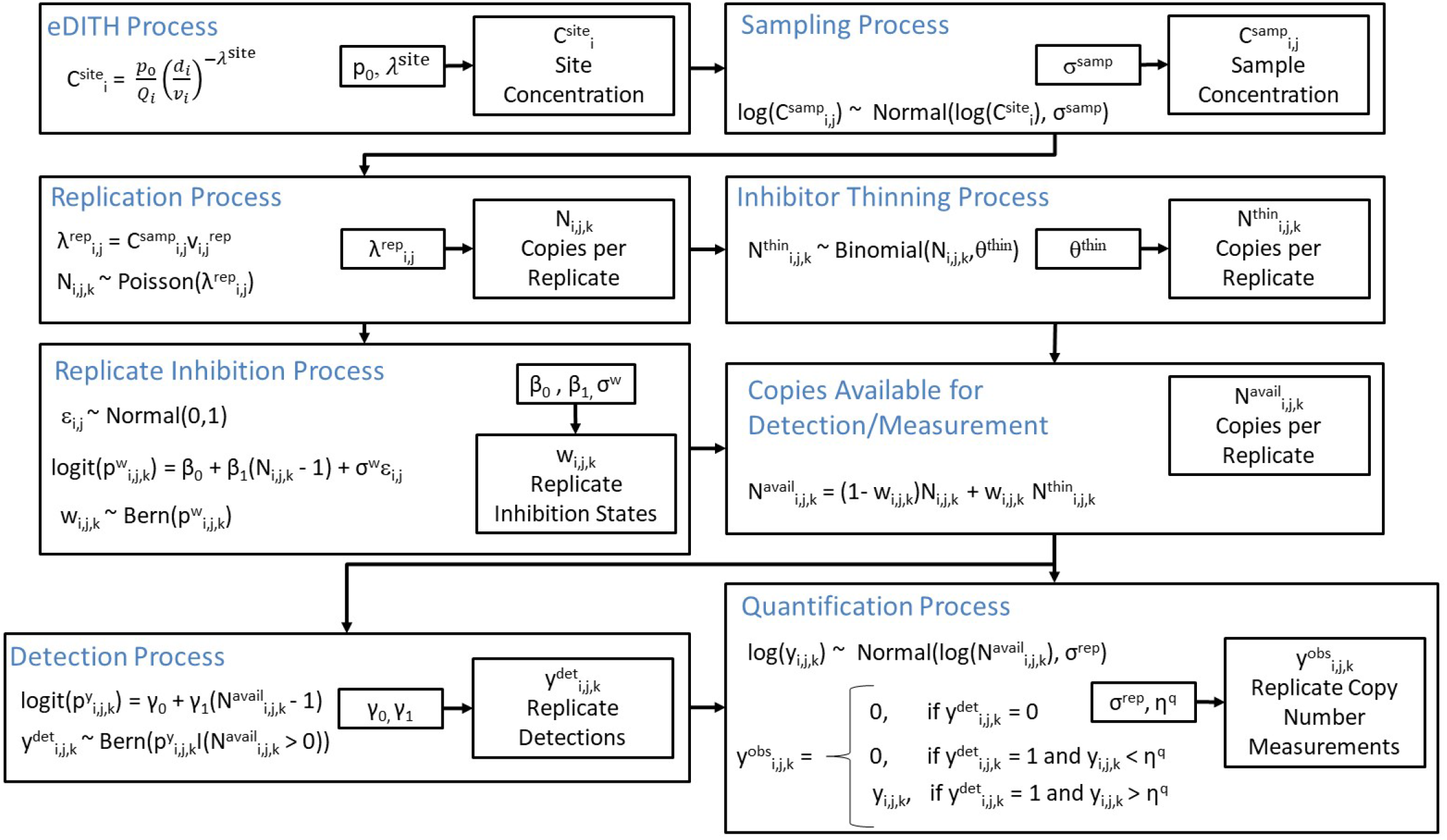
Model diagram with inhibitor processes. This diagram matches that in Figure 1, except between the replication and detection processes, there are 1) an inhibitor thinning process describing how many copies are available for detection and measurement for inhibited replicates, 2) a replicate inhibition process describing how replicates are inhibited, and 3) a description of how these two processes are combined to determine how many copies are available to be measured.

### Parameter Estimation

We estimated our model parameters using Markov chain Monte Carlo (MCMC) in the Nimble software (Version 1.0.1; de Valpine et al., 2017) in program R (Version 4.0.5; R Core Team, 2021). We used Nimble defaults for all MCMC sampler assignments, with some exceptions, specifically, 1) for the inhibitor models, we added user-defined samplers that were required to adequately sample the posterior (described in Appendix A), 2) we used a block Metropolis-Hastings update (Ponisio et al., 2020) for the eDITH model parameters (*p*_0_, *θ*^*site*^, *τ* or *λ*^*site*^) due to posterior correlation, and 3) we used a block Metropolis-Hastings update for 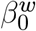 and 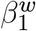 in the inhibitor models, also due to posterior correlation. Our priors can be found in Appendix A.

### Data Analysis

#### Model Comparisons

To investigate the evidence for and effects of both a non-constant eDNA removal rate with respect to time and eDNA inhibitors, we fit 4 models:

- Model I-PL – accounts for inhibitors, power law removal
- Model I-E – accounts for inhibitors, exponential removal
- Model N-PL – ignores inhibitors (null), power law removal
- Model N-E – ignores inhibitors (null), exponential removal

While the following are not completely conclusive on their own, we suggest that evidence supporting the inhibitor models would be 1) a strong lack of fit when ignoring inhibitors that is reduced when they are modeled (e.g., Model N-PL vs. Model I-PL), 2) a clear negative relationship between the probability of replicate-level undermeasurement events (hypothesized to be caused by inhibition) and the number of copies in a replicate, and 3) a positive relationship between the probability of replicate-level undermeasurement events and the replicate ΔCq.

If the inhibitor models are a good approximation of reality, we expected that when ignoring inhibitors, we will 1) underestimate site concentrations, with larger underestimation as site concentration decreases, 2) overestimate the eDNA removal rate due to under-measuring more at lower concentration sites further from the source, 3) overestimate both the sampling and measurement variability which must accommodate the effects of inhibition, and 4) underestimate the effect of replicate copy number on detection probability because the copy number in each replicate will be underestimated, on average. To compare the inhibitor models to the null models, we used posterior predictive checks (Gelman et al., 1996; Conn et al., 2018) and the Watanabe-Akaike Information Criterion (WAIC; Watanabe and Opper, 2010; Gelman et al., 2014). For the posterior predictive check, we used the observation model deviance as the discrepancy function (King et al., 2009; Conn et al., 2018). For the null models, the observation model deviance is

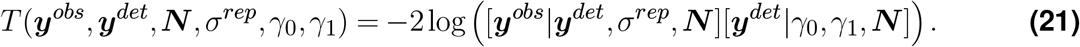

The observation model deviance for the inhibitor model is the same as for the null models, except we replace ***N*** with ***N***^*avail*^. For each data point, we computed the Bayesian P-value, the probability each observed data point’s discrepancy is more extreme than that of data simulated from the posterior. We considered observations with Bayesian P-values less than 0.05 to be poorly explained by the model and assumed models with more extreme data were less supported. We then used conditional WAIC to compare each model’s ability to predict data from these sites and samples (we were unable to compute a marginal WAIC to compare population-level predictive ability, i.e., new sites and samples). To assess the evidence for the hypothesis that the eDNA removal rate is not constant as a function of time, we used a less formal graphical assessment (Gelman et al., 1996) of the sampling process residuals, 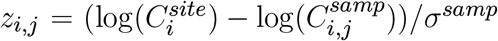, which are a pivotal quantity with a standard normal distribution (Conn et al., 2018). Here, we looked for deviations from normality and residual temporal/spatial correlation due to misspecification of the functional form of the removal model, a function of time that is converted to distance (space).

#### MCMC Details

For each of the 4 models, we ran 3 MCMC chains for 300,000 iterations each, with model parameters and latent variables thinned by 25 and 250 iterations, respectively. We discarded 25,000 pre-thinned iterations and assessed convergence using the Gelman-Rubin Statistic (Brooks and Gelman, 1998) ensuring the 95% CI upper bound was less than 1.1 for all parameters. Posterior modes were used as point estimates and 95% highest posterior density (HPD) intervals were used as interval estimates. Because the posteriors for ***N***, ***N*** ^*thin*^, ***N*** ^*avail*^, and ***C***^*sample*^ were often multimodal in the inhibitor models, we used the HDinterval R package (Meredith and Kruschke, 2020) to produce discontinuous HPD intervals.

### Simulation Study A

We conducted a simulation study to characterize how well parameters of each model are estimated in terms of relative bias, 95% coverage, and the coefficient of variation (CV) when using data similar to ours in terms of the data dimensions, hydrology, and parameter values. We simulated 100 data sets from each of four models using the parameter estimates from the field data, with the same number of sites, site distances, replicate number, and hydrological parameters. The simulation scenarios differed from the field data models in two ways–we excluded the thinning process at the first site due to the “plume effect” to assess how well the eDNA production rate, *p*_0_, can be estimated when ignoring this nuisance parameter, and we excluded the ΔCq covariate as it may not always be available.

For each simulated data set from the null and inhibitor models, we ran 3 MCMC chains for 150,000 and 300,000 iterations, respectively, and thinned posteriors by 25 to reduce file sizes. For null and inhibitor models, we discarded a minimum burn in of 5,000 and 35,000 pre-thinned iterations, respectively. After discarding this burn in, we computed the Gelman-Rubin statistic (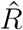 ; Gelman and Rubin, 1992) for each parameter, ensuring that the 95% confidence interval upper bound of the statistic was below 1.1. For posteriors with 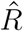 95% confidence intervals greater than 1.1 for any parameter, we discarded more burn in after visual inspection to meet this condition. For point estimates, we used the posterior mode, and for interval estimates, we used the 95% HPD interval. For each data set, we computed the CV for all parameters by dividing the posterior standard deviation by the absolute value of the posterior mode and we report the CV averaged across data sets for each model. One exception was for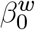, which had point estimates very close to zero, leading to large CVs which were not informative of parameter precision (Kvålseth, 2017). For this parameter, we computed the CV as the averaged standard deviation divided by the averaged posterior model.

### Simulation Study B

We conducted a second simulation study to demonstrate the effects of ignoring the replication and quantification process variation by using the mean concentration across replicates of a sample as observed data. We considered that sample concentrations could be estimated on the real scale or the log scale followed by a back-transformation to the real scale. This simulation study was targeted to show that 1) if quantification process variation is lognormal, bias is introduced to the sample concentration when averaging on the real scale, and 2) when replication and quantification process uncertainty is not partitioned, they inflate the sampling process standard deviation, *σ*^*samp*^. We expected the magnitude of these effects to vary by the number of replicates and the magnitude of the quantification process standard deviation, *σ*^*rep*^ (replication process variation is typically small relative to *σ*^*rep*^ and is determined by the Poisson mean). We considered 12 scenarios, using all combinations of *K ∈* (1, 3, 5) and *σ*^*rep*^ *∈* (0, 0.29, 0.68, 1.36). We chose *σ*^*rep*^ *∈* (0.29, 0.68) because these were the point estimates from the models that did and did not model the inhibition process, respectively. Then, we added a scenario of 0 where only the replication process variance is ignored and 1.36 to consider twice as much quantification variability than our highest estimate.

For each scenario, we simulated one data set from Model N-PL using *I* = 10 sites at distances that match our field sites up to a distance of 300 m. We set eDITH process parameters to have higher concentrations than our field data, did not thin the first site, and assumed perfect detection of copies in replicates in order to avoid simulating replicates with 0 copies where the log is undefined and this strategy of estimating the sample mean fails Chik et al. (2018). To achieve this, we set *p*_0_ = 30000, *λ*^*site*^ = 0.55, *θ*^*site*^ = 1, *γ*_0_ = 100 and *γ*_1_ = 0. We used the same hydrological measurements as in the analyses of field data, and set *η*^*q*^ = 0 so there is no data censoring at low concentration. For each data set corresponding to one scenario, we simulated *J* = 10000 samples per site with *σ*^*samp*^ = 1, similar to what we estimate in the inhibitor models. Note that there is no stochasticity in the eDITH process, so simulating many samples from a single data set is equivalent to simulating fewer samples from multiple data sets. For each scenario, we estimated sample means by averaging the sample replicates on the real and log scales. We then computed the mean and sample standard deviation of each set of sample mean estimates and computed their bias relative to estimates from the simulated data.

## Results

### Observed Data and Inhibition Assessment

We observed no amplification in negative controls or in samples collected upstream of the introduction point and positive controls amplified as expected. We observed substantially higher detection and copy number measurements using the short amplicon compared to the long amplicon assay. For the long amplicon assay, 37.5% of replicates were nondetections and 32.1% of samples failed to detect eDNA across all 5 replicates. The replicate-level quantitative data ranged from 1.71 - 4.90 log_10_ copies per liter and -0.29 - 2.90 log_10_ copies per reaction. Due to the poorer performance of the long amplicon assay, we used the short amplicon assay data for analysis.

The short amplicon replicate-level quantitative data ranged from 2.22 - 5.91 log_10_ copies per liter and 0.22 - 3.91 log_10_ copies per reaction (Figure 3a). We observed 24 replicates (out of 390) that failed to detect eDNA, with 1, 5, 7, 1, 2 and 8 failed detections at distances of 0.05, 40, 300, 400, 500, and 1000 m, respectively. Three samples failed to detect eDNA across all five replicates, one sample each at distances of 40, 300, and 1000 m. Our inhibition assessment did not indicate inhibition for any replicates using the decision rule of Δ*Cq ≥* 3. Shifts in Cq ranged from -0.13 to 1.60 with a mean of 0.11 cycles. The distribution of ΔCq was right skewed, with most values clustered slightly above zero (Figure 3b).

**Figure 3.**
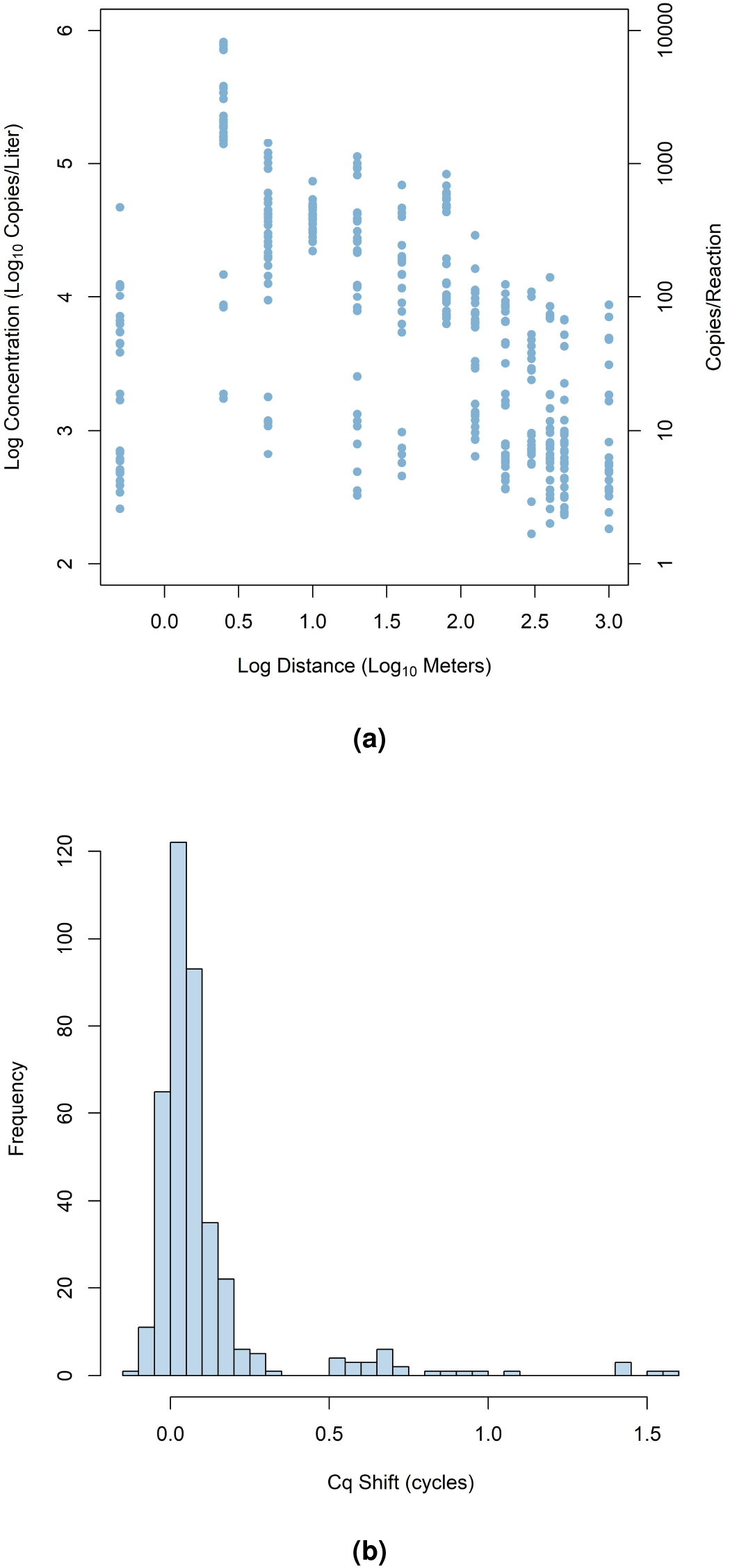
Observed replicate-level quantitative measurements (a) and Cq shifts (b). (a) Replicate-level log_10_ concentrations (left y-axis) and copies per reaction (right y-axis) as a function of log_10_ distance from the release location. (b) Distribution of replicate-level Cq shifts between internal positive controls and plate-level mean Cq for uninhibited control samples.

### Analysis of Field Data

The four models we considered produced substantially different parameter estimates, leading to different inferences about the ecological and observational processes that produced our data (Table 1). The inhibitor models estimated that the sampling process standard deviation, *σ*^*samp*^ was smaller and the quantification process standard deviation, *σ*^*rep*^ was larger, compared to the null models. The inhibitor models further estimated that the removal rate (governed by *λ*^*site*^ or *τ*) was lower compared to the null models, and that site concentrations were higher, particularly at the sites farthest from the source (Figure 4). The power law removal models estimated the eDNA production rate at the source to be much larger than the exponential removal models (Figure 5) and correspondingly, estimated the percent of total concentration that is measurable at the first site, *θ*^*site*^ to be lower (Table 1). Finally, the effect size of replicate copy number on replicate detection probability was estimated to be greater in the inhibitor models relative to the null models.

**Table 1.** Parameter point estimates for all 4 models applied to the field data. Model I-PL accommodates inhibitors and has power law removal, Model I-E accommodates inhibitors and has exponential removal, Model N-PL does not accommodate inhibitors and has power law removal, Model N-E does not accommodate inhibitors and has exponential removal.

| Process | Par | Model |  |  |  |
| --- | --- | --- | --- | --- | --- |
|  |  | I-PL | I-E | N-PL | N-E |
| eDITH | $p_0$ | 5914.94 | 414.66 | 11486.98 | 257.53 |
| eDITH | $\lambda^{site}$ | 0.58 | . | 0.89 | . |
| eDITH | $\tau$ | . | 1425.53 | . | 867.18 |
| eDITH | $\theta^{site}$ | 0.03 | 0.34 | 0.01 | 0.16 |
| Sampling | $\sigma^{samp}$ | 0.91 | 1.20 | 1.34 | 1.70 |
| Replicate Inhibition | $\beta_0^w$ | 1.47 | 1.32 | . | . |
| Replicate Inhibition | $\beta_1^w$ | -0.02 | -0.02 | . | . |
| Replicate Inhibition | $\beta_2^w$ | 1.18 | 1.16 | . | . |
| Replicate Inhibition | $\sigma^w$ | 3.28 | 3.37 | . | . |
| Inhibitor Thinning | $\theta_0$ | -2.31 | -2.34 | . | . |
| Detection | $\gamma_0$ | -2.60 | -2.45 | -2.24 | -2.05 |
| Detection | $\gamma_1$ | 2.72 | 2.69 | 1.26 | 1.25 |
| Quantification | $\sigma^{rep}$ | 0.29 | 0.29 | 0.68 | 0.69 |

**Figure 4.**
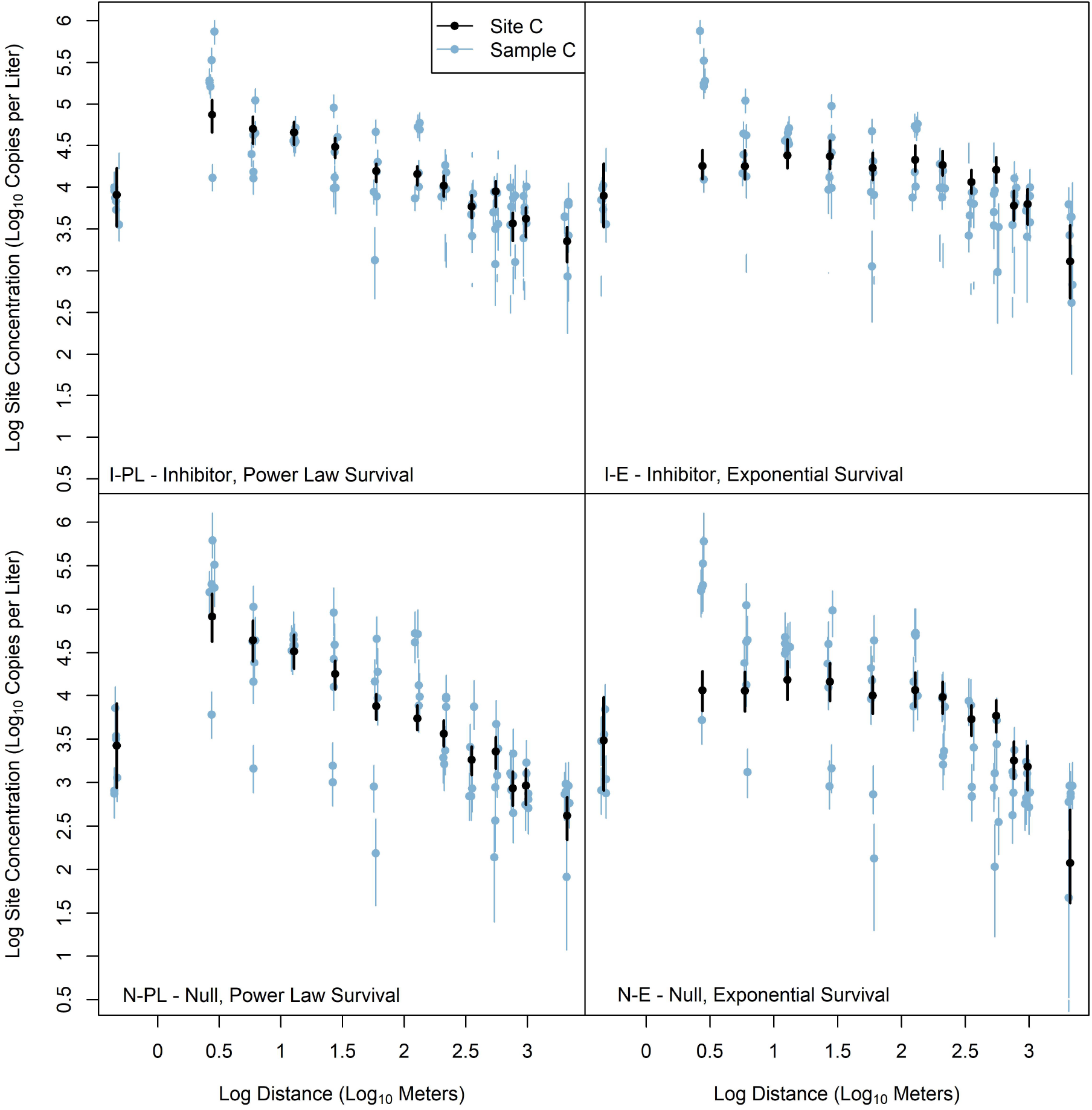
Posterior point and interval estimates of site and sample concentration as a function of distance for all four models. Note, some 95% HPD intervals are discontinuous due to multimodal posterior distributions. Estimates from Models I-PL and I-E accounting for inhibitors are generally higher, and to a greater magnitude as distance increases. Estimates from Models I-PL and N-PL are higher at sites 2-4 compared to Models I-PL and N-PL, respectively, due to an increasing removal rate through time.

**Figure 5.**
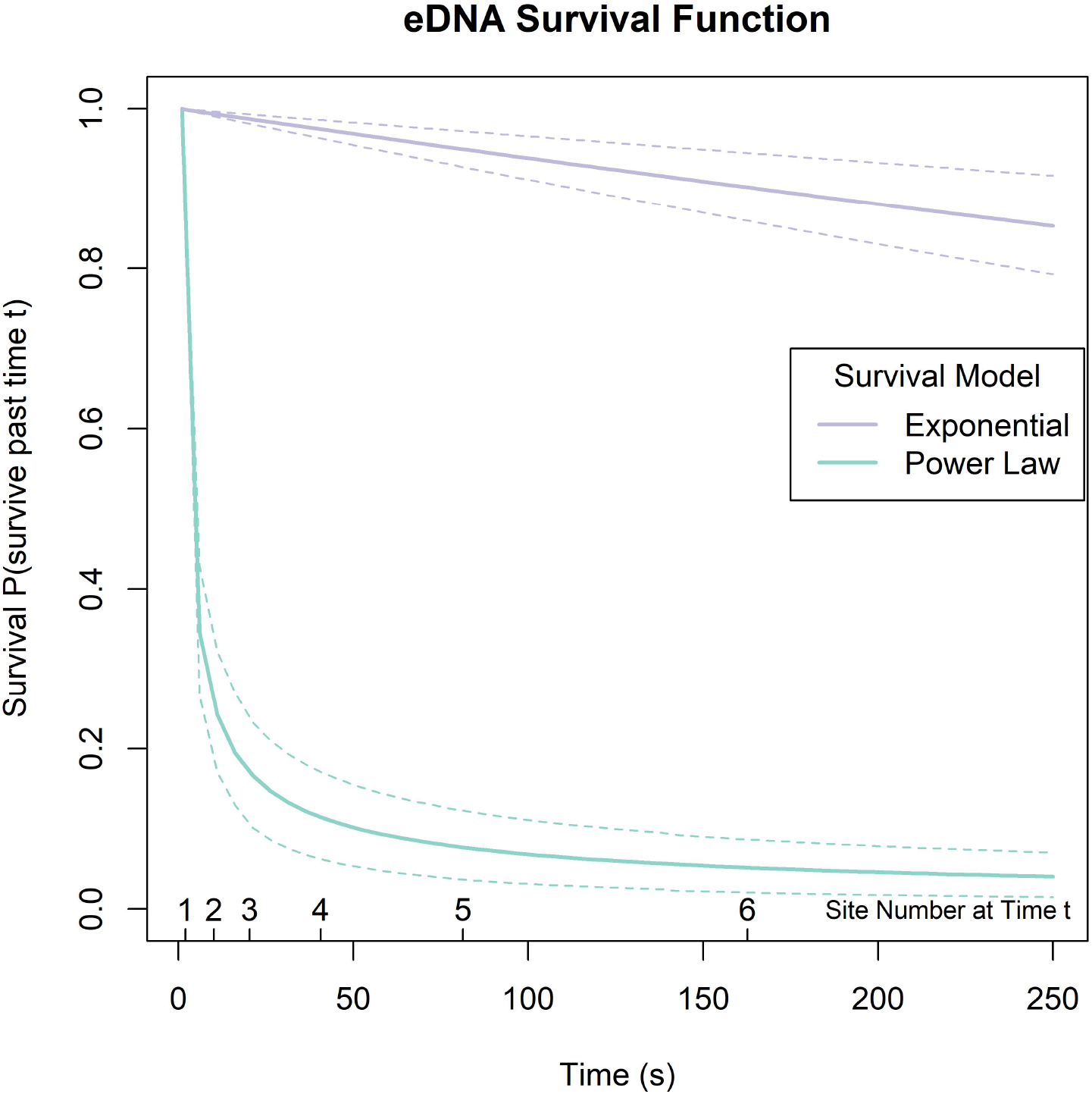
Survival and Log Hazard functions from Models I-PL and I-E, with power law and exponential survival, respectively. Under the power law model, there is more eDNA estimated to be added at the source location relative to the exponential, and faster removal that slows down through time. The expected time at which eDNA reaches sites 1-6 are indicated on the plots, which were derived from our mean velocity measurement.

The Bayesian p-values from the deviance discrepancy function identified 14 samples as having replicates that were more extreme than expected under both versions of the null models (Table 2). These samples were characterized by having lower copy numbers, with one or more replicate measurements being much larger than the others, and they disproportionately came from the sites with the lowest measurable concentrations (first site with plume effect and sites farthest from the source). For the inhibitor models, the Bayesian p-values identified 4 samples having replicate measurements that were more extreme than expected by the model. These replicates were also imbalanced across replicates but to a lesser or greater degree than expected compared to our thinning rate estimate, inv. logit(*θ*_0_) = 0.09, suggesting heterogeneity in the sample-level thinning rate. WAIC favored the inhibitor models over the null models, with the increased number of effective parameters outweighed by the substantially higher log posterior predictive density (Table 3). Plots of the sampling process residuals from the inhibitor models (Figure 6) suggested that the exponential removal model (Model I-E) was less consistent with the assumption of normality on the log scale, with more of a quadratic pattern in the residuals plotted vs. distance from source (ignoring the thinned first site). A Shaprio-Wilk test for normality of the residual point estimates rejected the null hypothesis of normality for Model I-E (p=0.005), but not Model I-PL (p=0.279). Conversely, WAIC favored Model I-E over I-PL by 1.12 units.

**Table 2.** Samples with replicates where Bayesian p-values from posterior predictive checks indicated lack of fit, defined as the p-value being less than 0.05. Samples are listed by model (“Null” and “Inhibit” without and with modeling inhibitors, respectively), site, and sample, along with the observed data for each sample replicate. Null models are N-E and N-PL and inhibitor models are I-E and I-PL.

| Model | Site | Sample | Rep 1 | Rep 2 | Rep 3 | Rep 4 | Rep 5 |
| --- | --- | --- | --- | --- | --- | --- | --- |
| Null | 1 | 1 | 2.57 | 66.33 | 62.70 | 3.41 | 3.86 |
| Null | 1 | 3 | 0 | 7.08 | 4.88 | 472.19 | 4.16 |
| Null | 1 | 6 | 125.24 | 71.73 | 4.75 | 121.01 | 5.86 |
| Null | 8 | 5 | 62.79 | 6.35 | 164.17 | 11.97 | 8.67 |
| Null | 10 | 1 | 110.24 | 5.66 | 9.56 | 7.31 | 5.53 |
| Null | 11 | 3 | 8.16 | 74.08 | 7.98 | 10.39 | 141.60 |
| Null | 11 | 4 | 68.67 | 7.37 | 7.88 | 11.77 | 6.02 |
| Null | 11 | 6 | 70.95 | 2.56 | 3.30 | 84.97 | 4.21 |
| Null | 12 | 1 | 2.31 | 7.94 | 5.57 | 2.64 | 52.09 |
| Null | 12 | 2 | 8.30 | 9.61 | 4.28 | 66.94 | 9.94 |
| Null | 12 | 3 | 42.74 | 5.61 | 0 | 3.24 | 2.38 |
| Null | 13 | 3 | 2.42 | 48.81 | 18.51 | 1.83 | 0 |
| Null | 13 | 4 | 31.04 | 3.61 | 0 | 87.01 | 4.80 |
| Null | 13 | 5 | 70.73 | 3.49 | 5.00 | 8.21 | 5.42 |
| Inhibit | 1 | 3 | 0 | 7.08 | 4.88 | 472.19 | 4.16 |
| Inhibit | 1 | 4 | 44.27 | 102.19 | 38.40 | 16.92 | 18.84 |
| Inhibit | 6 | 4 | 202.90 | 54.30 | 62.58 | 90.22 | 77.76 |
| Inhibit | 8 | 4 | 67.78 | 74.98 | 76.75 | 291.60 | 76.44 |

**Table 3.**
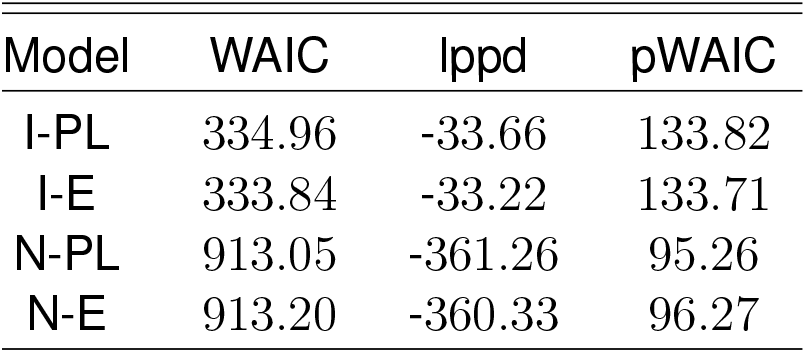
WAIC Table showing the WAIC, log posterior predictive density (lppd) and effective number of parameters (pWAIC) for all 4 models.

**Figure 6.**
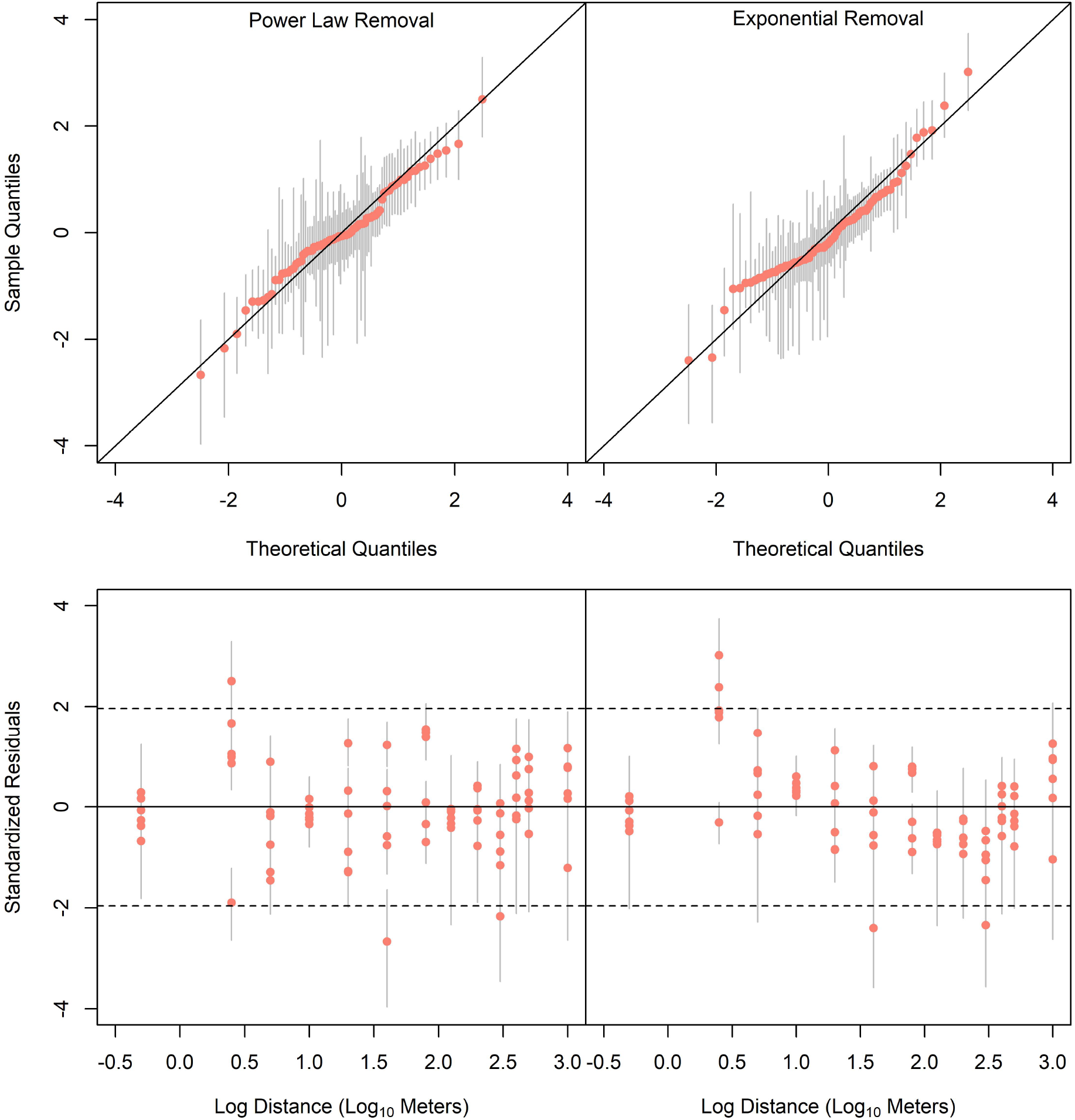
Plots of the log sample concentration standardized residuals with 95% HPD intervals from Models I-PL and I-E assuming eDNA removal follows a power law (left) or exponential (right) model. On top are quantile-quantile plots comparing sample quantiles to theoretical quantiles where a 1:1 relationship indicates consistency with normality. On bottom are standardized residuals plotted against log distance from the release location where residuals should have a mean of zero and standard deviation of 1 across all distances if the removal model is correctly specified. The dotted lines indicate the 2.5 and 97.5% quantiles of Normal(0,1) distribution.

Parameter estimates with posterior standard deviations and 95% HPD intervals for Model I-PL can be found in Table 4. Model I-PL estimated that many replicates in the range of 1.5 - 2.5 log_10_ copies were substantially undermeasured, but after accounting for the inhibition process, the copies available to be measured corresponded well with the measured copy numbers (Figure 7). In Model I-PL, the probability of inhibition was estimated to be inv. 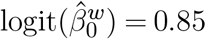 for a replicate with 1 copy (and random effect value set to 0), with a negligible probability of inhibition over 300 copies per replicate (Figure 8). The ΔCq covariate was estimated to be positively-related to the probability of inhibition (Table 4), with a 95% HPD that did not overlap 0 and a posterior probability of being greater than 0 equal to 0.99.

**Table 4.** Parameter point and interval estimates and posterior standard deviations from Model I-PL that accommodates inhibitors and considers power law removal.

| Process | Par | Est | SD | Lower | Upper |
| --- | --- | --- | --- | --- | --- |
| eDITH | $p_0$ | 5914.94 | 2853.66 | 2758.90 | 12983.98 |
| eDITH | $\lambda^{site}$ | 0.58 | 0.07 | 0.46 | 0.73 |
| eDITH | $\theta^{site}$ | 0.03 | 0.03 | 0.01 | 0.10 |
| Sampling | $\sigma^{samp}$ | 0.91 | 0.10 | 0.75 | 1.12 |
| Replicate Inhibition | $\beta_0^w$ | 1.47 | 0.98 | -0.19 | 3.64 |
| Replicate Inhibition | $\beta_1^w$ | -0.02 | 0.01 | -0.05 | -0.01 |
| Replicate Inhibition | $\beta_2^w$ | 1.18 | 0.54 | 0.18 | 2.32 |
| Replicate Inhibition | $\sigma^w$ | 3.28 | 0.78 | 2.18 | 5.11 |
| Inhibitor Thinning | $\theta_0$ | -2.31 | 0.11 | -2.53 | -2.11 |
| Detection | $\gamma_0$ | -2.60 | 1.31 | -5.74 | -0.78 |
| Detection | $\gamma_1$ | 2.72 | 1.78 | 1.14 | 7.18 |
| Quantification | $\sigma^{rep}$ | 0.29 | 0.02 | 0.26 | 0.33 |

**Figure 7.**
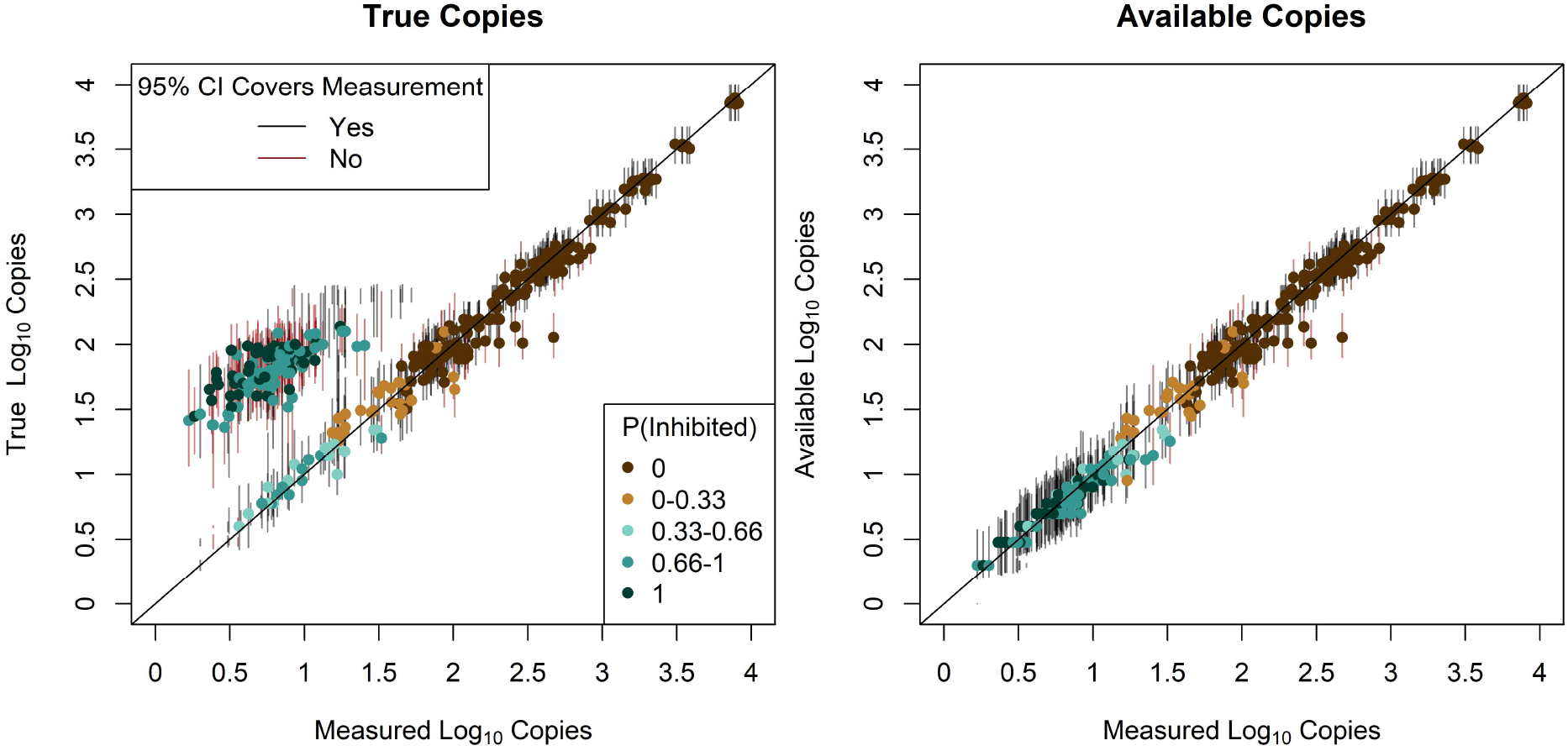
Comparison of the relationship between measured and true copies in each replicate (left) and measured and available copies in each replicate (right) from Model I-PL accounting for inhibitors. In each plot, the posterior probability of inhibition is indicated by the color of the point estimate and 95% HPD intervals are depicted, some of which are discontinuous due to multimodal posteriors. In the plot on the left, the group of true copies measured too low with poor coverage correspond to the samples to which the model assigns a high posterior probability of inhibition. After accounting for inhibition (plot on right), the relationship between measured and available copies is roughly linear with coverage 0.87.

**Figure 8.**
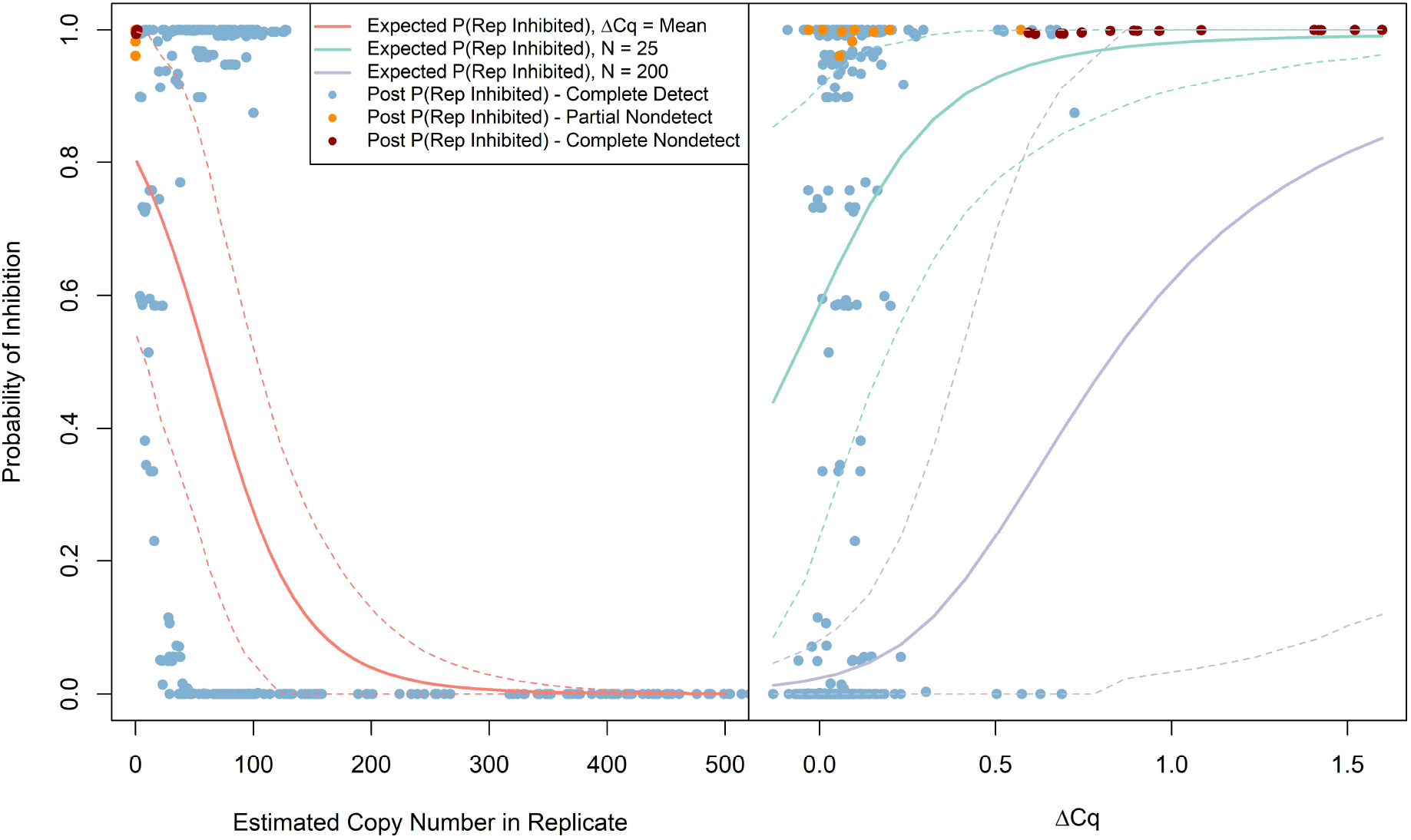
Plots depicting the effects of copy number and ΔCq on the expected probability of inhibition (posterior mean and 95% HPDs),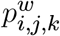, and the posterior probability of inhibition, P(*w*_*i,j,k*_ = 1). Replicates from samples with no detections (complete nondetects) are colored red, and replicates without detections in samples where other replicates were detected (partial nondetects) are colored orange. The probability of inhibition declines as a function of replicate copy number, with complete and partial nondetects all being estimated to have very few copies (left). The probability of inhibition increases with ΔCq, particularly for replicates with fewer copies (N=25 scenario)

### Simulation Study A

Our ability to estimate parameters in terms of bias, coverage, and precision (CV) varied across parameters and models, with generally better estimates from null models over inhibitor models and exponential over power law removal models (Table 5). For the null models, coverage was roughly nominal, but the detection parameters were moderately biased (−6.45 - 16.4%) and estimated imprecisely (CVs of 62.4 - 144.3%). Further, the null model with exponential removal estimated the eDNA production rate, *p*_0_ with minimal bias (2.6%) and a CV of 29.0%, whereas the estimate from the null model was positively biased by 15%, and less precise (CV of 52.8%). The sampling and measurement standard deviations, *σ*^*samp*^ and *σ*^*rep*^, were estimated with minimal bias (<2%) in the null models.

**Table 5.** Relative bias, 95% coverage, and coefficient of variation of model parameters in the simulation study for the four models considered. “I” indicates “inhibitor,” “N” indicates “Null,” “PL” indicates “power law” removal, and “E” indicates “exponential” removal. “Bias” is the estimated relative bias as a percentage, “Cover” is the estimated 95% coverage of 95% HPD intervals, and “CV” is the average coefficient of variation across simulated data sets as a percentage. Note, for 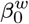 which had parameter estimates near 0, the CV was calculated using the point estimate and standard deviation averaged across data sets.

| Par | I-PL |  |  | I-E |  |  | N-PL |  |  | N-E |  |  |
| --- | --- | --- | --- | --- | --- | --- | --- | --- | --- | --- | --- | --- |
|  | Bias | Cov | CV | Bias | Cov | CV | Bias | Cov | CV | Bias | Cov | CV |
| $p_0$ | 13.4 | 0.93 | 36.6 | -1.8 | 0.97 | 20.1 | 15.1 | 0.94 | 52.8 | 2.6 | 0.95 | 29.0 |
| $\lambda^{site}$ | 5.9 | 0.90 | 9.5 | . | . | . | 3.6 | 0.91 | 7.9 | . | . | . |
| $\tau$ | . | . | . | 5.7 | 0.93 | 31.3 | . | . | . | -3.5 | 0.95 | 22.7 |
| $\sigma^{samp}$ | 4.9 | 0.91 | 10.8 | 1.9 | 0.92 | 11.0 | 1.5 | 0.93 | 9.6 | 1.1 | 0.92 | 9.3 |
| $\beta_0^w$ | -34.8 | 0.89 | 53.5 | -33.7 | 0.94 | 61.4 | . | . | . | . | . | . |
| $\beta_1^w$ | 10.1 | 0.92 | 55.0 | 10.7 | 0.95 | 32.6 | . | . | . | . | . | . |
| $\sigma^w$ | -11.2 | 0.94 | 24.0 | -14.7 | 0.93 | 23.5 | . | . | . | . | . | . |
| $\theta_0$ | 1.5 | 0.93 | 4.7 | 0.2 | 0.95 | 4.28 | . | . | . | . | . | . |
| $\gamma_0$ | 22.4 | 0.95 | 52.3 | 22.1 | 0.95 | 54.2 | 11.3 | 1.00 | 62.4 | 16.4 | 0.97 | 81.1 |
| $\gamma_1$ | -10.9 | 0.97 | 85.9 | -7.1 | 0.99 | 90.1 | -6.5 | 0.99 | 97.0 | 1.9 | 1.00 | 144.3 |
| $\sigma^{rep}$ | -0.5 | 0.96 | 5.6 | -0.4 | 0.96 | 5.4 | -0.1 | 0.95 | 4.6 | -0.5 | 0.95 | 4.5 |

For the inhibitor models, we saw results similar to the null models. Overall, coverage was roughly nominal for the inhibitor models with the possible exception of reduced coverage for some parameters in model I-PL (our estimates are subject to some sampling variability with only 100 data sets). The detection parameters were biased (−10.9 - 22.4%) and estimated imprecisely (CVs of 52.3 - 90.1%). As in the null models, the eDNA production rate, *p*_0_, was estimated with minimal bias in model I-E with exponential removal (−1.8%) and positive bias in model I-PL with power law removal (13.4%). The *p*_0_ estimates in the inhibitor models were more precise as judged by the CV, likely due to the larger simulated values used in the null models leading to larger posterior modes. All parameters determining the probability of inhibition were estimated with bias, particularly the intercept 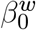, which was negatively biased by roughly 34% in both inhibitor models. As in the null models, the sampling and measurement standard deviations, *σ*^*samp*^ and *σ*^*rep*^, were estimated with minimal bias (<2%), except for *σ*^*samp*^ in model I-PL with a bias of 5.9%.

### Simulation Study B

When producing sample mean estimates by taking the mean of replicates on the log scale, bias in estimated sample means was negligible, whereas taking the mean of replicates on the real scale introduced bias as the sampling process standard deviation, *σ*^*rep*^, and the number of replicates, *K*, increased (Table 6). This bias was negligible for the levels of *σ*^*rep*^ we estimated in the null and inhibitor models for our field data. For both averaging methods, bias in *σ*^*rep*^ estimates increased with *σ*^*rep*^, decreased with *K*, and was negligible for our inhibitor model *σ*^*rep*^ estimate except for *K* = 1. For our null model *σ*^*rep*^ estimate, bias was 5.4% with the maximum number of replicates we considered (*K* = 5).

**Table 6.** Relative bias in estimated sample concentrations (Mean Bias) and Sampling Process standard deviation, *σ*^*samp*^, (SD Bias) from simulated data sets when estimating sample means by averaging replicates on the real vs. log scale. Scenarios vary *σ*^*rep*^ and *K*, each using 1 simulated data set across 10 sites with 10000 samples per site. For all scenarios, the sampling process variance was *σ*^*samp*^ = 1. Averaging replicates on the log scale produces unbiased estimates, while averaging on the real scale introduces positive bias as *σ*^*rep*^ and *K* increase (note, when K=1, there is no averaging across replicates). Ignoring the variation due to the Replication and Quantification Processes increasingly introduces bias into the Sampling Process standard deviation as *σ*^*rep*^ rises and *K* declines.

| $\sigma^{rep}$ | K | Mean Bias (%) | | SD Bias (%) | |
| --- | --- | --- | --- | --- | --- |
|  |  | Real | Log | Real | Log |
| 0 | 1 | -0.01 | -0.01 | 0.19 | 0.19 |
| 0 | 3 | 0.00 | -0.01 | 0.07 | 0.14 |
| 0 | 5 | 0.00 | -0.01 | 0.05 | 0.13 |
| 0.29 | 1 | 0.01 | 0.01 | 4.28 | 4.28 |
| 0.29 | 3 | 0.23 | -0.01 | 1.47 | 1.51 |
| 0.29 | 5 | 0.29 | -0.01 | 0.92 | 0.96 |
| 0.68 | 1 | 0.02 | 0.02 | 21.05 | 21.05 |
| 0.68 | 3 | 1.22 | -0.01 | 8.67 | 7.83 |
| 0.68 | 5 | 1.51 | -0.01 | 5.30 | 4.62 |
| 1.36 | 1 | 0.06 | 0.06 | 69.25 | 69.25 |
| 1.36 | 3 | 4.27 | 0.00 | 34.50 | 27.44 |
| 1.36 | 5 | 5.43 | 0.00 | 24.48 | 17.06 |

## Discussion

We developed a hierarchical model for eDNA fate and transport experiments that accommodates more mechanistic detail about how eDNA is detected and measured than existing models. We also conducted an eDNA release experiment to demonstrate the utility of this modeling approach for estimating eDNA removal parameters in the presence of ecological and measurement variability. Two distinctive features of the model are that site concentrations are modeled as the product of an eDNA removal process as a function of time (using the eDITH approach of Carraro et al., 2018) and that replicate copy numbers are regarded as latent variables modeled with a count distribution. The latter feature has several advantages for low concentration samples and facilitates modeling detection, measurement error, and sources of bias as functions of the ecological quantity being measured–the discrete number of copies in each replicate. Using this model, we were able to provide evidence that the eDNA removal rate in our experiment declined through time and that the allocation of measurable copies across replicates was overdispersed relative to the Poisson in a subset of samples. We then developed a model for how eDNA inhibitors might reduce the measurable copy numbers in the replicates and show the pattern of overdispersion in our observed data is largely consistent with this model.

Treating copy numbers in replicates as count random variables provides several advantages when site and sample concentrations are low. To date, quantitative eDNA analyses have typically assumed sample or replicate concentrations are normally-distributed, usually on the log scale (Carraro et al., 2018; Espe et al., 2022; Shelton et al., 2019). Our model highlights that replicate-level measurement error is the product of both sampling variation when allocating copies to the replicates (Dube et al., 2008; Lesperance et al., 2021) and the measurement error conditioned on the number of copies in each replicate (due to variation in factors such as reading fractional cycle numbers; Shelton et al., 2019). An implication of a count model for the replication process is that a normal approximation will be a less appropriate when measuring small copy numbers (Espe et al., 2022) that are typical of many eDNA applications. Then, when sample concentrations are low enough that qPCR replicate-level zeros are observed, it is unknown whether a zero is observed because there were no target DNA copies allocated to a replicate or because the allocated copies were undetected (Dube et al., 2008; Lesperance et al., 2021). To compute an unbiased sample mean concentration in the presence of observed zeros, the replication process zeros need to be included, as do the true allocated copy numbers for replicates with failed detections, which are not observed.

Techniques that allow a probabilistic interpretation of zeros and non-detections have been recognized as critical by many disciplines, like analytical chemistry and public health (e.g., Chik et al., 2018), and have been used by eDNA practitioners when computing the limit of detection in qPCR (Dube et al., 2008; Lesperance et al., 2021). By conditioning detection on copies being present in replicates, and relating the detection probability to the number of copies in a replicate, our model can probabilistically resolve these sources of zeros and produce less biased estimates of replicate copy numbers and sample concentrations in this situation. In practice, the accuracy of the resulting estimates will depend on how well model assumptions are met, and estimates may be very imprecise when samples have concentrations so low that most replicates are true sampling zeros. Further, in Simulation Study A, we show that the detection parameters that are a function of latent copy numbers are weakly identifiable, estimated with low precision and moderate bias. We assume this is due to the magnitude of uncertainty in the estimates of the latent copy numbers near zero, which is greater in the inhibitor models where the bias in the detection probability of one copy is larger. Therefore, the performance of this model in terms of bias and precision of the ecological parameters in scenarios with lower sample concentrations than we observed and/or greater sampling variability should be further investigated. This approach of partitioning variation between the allocation of copies and measurement of copies may also be used in standard curve experiments (e.g., Klymus et al., 2020) to relate Cq to the realized, instead of expected, copy number of standards. Doing so could reduce the nonlinearity seen in standard curve calibration regressions (Klymus et al., 2020) due to observing true and false zeros at low concentration standards.

Including latent replicate copy numbers in the model also provides a means for detecting deviations from Poisson variability in the replication process, and exploring hypotheses about potential causes of this assumption violation. We were able to use posterior predictive checks to identify lack of fit in our null models consistent with overdispersion in the number of copies allocated to replicates. This overdispersion was only seen in a subset of samples, and appeared stochastic in nature, with replicates tending to show a bimodal pattern of high and low measurements in affected samples. In theory, general overdispersion could result from sample-level variability in measurement error given the number of copies allocated to each replicate, but we were unable to devise a measurement error hypothesis that explained sample-level heterogeneity and the apparent bimodality in affected samples. Further, we did not find general overdispersion hypotheses, such as pipetting error, to be plausible because most samples were consistent with Poisson replication process variability.

A benefit of our hierarchical modeling approach is that we can construct and evaluate hypotheses for plausible mechanisms of this overdispersion pattern. Using the common decision rule of ΔCq≥ 3 (e.g., Hartman et al., 2005; McKee et al., 2015; Goldberg et al., 2016), none of our samples would be classified as inhibited (Figure 3b). However, given the general vulnerability of PCR quantification to inhibition (Opel et al., 2010; Sidstedt et al., 2020), the lack of a standard criterion to describe degrees of inhibition (Lance and Guan, 2020), and variation in how different assays respond to the same inhibitor (Lance and Guan, 2020), we suspected this decision rule may not be well calibrated for our assays and inhibition may have gone undetected using this method. We used our extended hierarchical model to evaluate the hypothesis that eDNA inhibitors were reducing detection probability and measured copy numbers in our field data, and these inhibitor effects were stochastic, leading to undermeasurement of some replicates of some samples. In the model, inhibitors have a continuous effect, causing undermeasurement, then failed detection, depending both on the starting copy number and the magnitude of the underestimation caused by the inhibitors.

We suggest that the posterior predictive checks provide support for the inhibitor models over the null models. The inhibitor model did not provide a clear indication of lack of fit for 13/14 of the poorly explained samples from the null model (Table 2), and the 4 samples that were identified by the posterior predictive check as indicating lack of fit in the inhibitor model were cases that are consistent with inhibition, but with a thinning rate lower or larger than our estimate of *θ*^*thin*^ = 0.09 (assuming binomial variability). This extra-binomial variability could be accommodated with a sample-level random effect in the inhibitor thinning process, or perhaps both the probability and magnitude of inhibition could be related to the same latent variable representing inhibitor concentration in a sample. Conditional WAIC also highly favored the inhibitor model, suggesting it is a better model to predict observed data from the samples we collected. A second line of evidence for the inhibitor hypothesis is that the probability of inhibition was positively associated with the ΔCq covariate with a posterior probability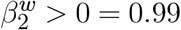. Still, we cannot conclusively rule out other causes of the overdispersion we saw in our data, and it may result from more than one cause.

Hypothetically interpreting our field data in light of the inhibitor model, inhibition started to occur in the range of 100-400 copies per replicate. As copy number approached zero, between 60 and 100% of the replicates were likely to be inhibited (interpreting 95% CIs in Figure 8). If inhibitors were as prevalent in our data as our model suggests, this would indicate that exogenous IPC Cq shifts may not always have high statistical power to detect inhibition at magnitudes that substantially affect measurements. Whether the source of overdispersion we detected was due to a mechanism increasing variability or one causing systematic undermeasurement, like inhibitors, is of high importance. The latter will cause us to systematically underestimate site and sample concentrations disproportionately such that as concentration declines, the rate of eDNA removal is overestimated and the sampling process variability (e.g., attribute measurement variability to ecological variability) is inflated. Our model provides a framework to compare alternative hypotheses, using either field or experimentally manipulated samples, to such extent that these hypotheses imply that different data will be observed. A final caveat is that the signal of inhibition is eroded as copy number approaches zero because of the magnitude of measurement error given the available copy number. For the inhibitor model parameters to be identifiable, we require samples with concentrations high enough that the magnitude of imbalance (relative to Poisson sampling variability) across replicates is large relative to measurement error, but not so large that inhibition did not occur. Lower concentration samples are more likely to have all replicates inhibited, so no imbalance is observable. Our simulation results indicate that the probability of inhibition parameters (***β***^*w*^ and *σ*^*w*^) were weakly identifiable for data sets similar to ours, and we expect they will be completely unidentifiable in many cases, such as our long amplicon assay results with high levels of nondetection.

A key advantage of hierarchical modeling is that variability can be estimated separately for individual processes, which allows for more informed hypothesis testing. Simulation Study B, for example, demonstrates that without a hierarchical framework the variability from the replication and quantification processes would be indistinguishable from sampling process variability. Such a conflation could easily be misinterpreted as ecological variability. The magnitude of measurement variability that is erroneously propagated to sampling variability depends on the number of replicates (averaging more numbers increases the accuracy of the mean) and the magnitude of the measurement error (Table 6). Using sample means computed in this manner as data in downstream analyses amounts to “doing Statistics on Statistics” (Royle and Dorazio, 2008) and is quite common in eDNA studies that scale up observations to the site (e.g., Laporte et al., 2020), but this will not alter ecological inference if the bias introduced is negligible. Given our estimates of sampling and measurement error, particularly from the inhibitor model, minimal bias is introduced to the sample mean concentration and standard deviation estimates when averaging replicates on the log scale to produce sample means. However, we show that, in principle, this practice conflates sources of variation, and the bias introduced can be large under certain scenarios. We also show that the sample mean concentrations are biased when averaging on the real scale instead of the log scale because we assume measurement error conditioned on copy number is lognormal, as we expect when errors (using qPCR) on the scale of Cq are normal. In addition to the challenges of discriminating sampling from detection zeros described above, sample means cannot be computed on the log scale when there are one or more zeros. Hierarchical modeling, generally, provides a means of separating measurement from ecological variability, and our model, specifically, provides a solution for incorporating the uncertainty in the source of zeros.

By using the eDITH model to describe eDNA removal as a function of time, we were able to provide evidence that the eDNA removal rate was not constant through time. Figure 6 shows the sampling process residuals better meet the normality assumption on the log scale under power law removal compared to exponential removal, with the latter showing more of a quadratic pattern in the residuals vs. distance. While difficult to quantify, we suggest this is evidence for misspecification of the functional form in the exponential model that is resolved by allowing the removal rate to decline with time. Conditional WAIC favored the exponential model by 1.1 unit, which we interpret as evidence that these models have similar predictive performance for our sites and samples. If the power law model is closer to the truth, the eDNA production rate parameter, *p*_0_, is challenging to estimate given our study design and the magnitude of sampling variance we encountered. In Simulation Study A, we see that *p*_0_ is estimated with negligible bias in the exponential removal models, but moderate positive bias in the power law removal models. Then, the CV is roughly twice as large for the power law removal models. We attribute this to the challenge of extrapolating the pattern in observed concentrations back to the source location when concentration is declining extremely rapidly (Figure 5). Using the exponential model if the power law model is more appropriate will negatively bias the eDNA production rate at the source, which will be propagated to metrics like site abundance. While not conclusive on its own, we suggest our study adds to the evidence for the hypothesis that, in the context of a mass-balance model of eDNA production, transport, and decay, the removal term is large and the transport term is small (Tillotson et al., 2018).

Finally, we discuss implications of our model and results for applications of the eDITH model to observational data and for occupancy analyses using detection data only. To our knowledge, the eDITH model has only been applied to observational data sets where: 1) the concentration observed at one site is the product of multiple upstream sites with different eDNA production rates and 2) release locations are estimated because they are not known. In contrast, our study system used a single, known release location. If eDNA decay declines at the rate we estimated, the majority of the total eDNA produced in a river network would be ignored, and the resulting site concentrations would be underestimated. Further, this underestimation would be predominately close to the sources, so measured site concentrations would not be related to true site concentrations by the same proportionality constant across space (inhibition could also cause deviations from a fixed proportionality constant). The assumptions of the eDITH model are hard to scrutinize given the number and magnitude of sources of ecological and measurement variability and the number of parameters and latent variables that must be estimated. We demonstrated how some assumptions can be better tested in more simple experiments and recommend more and better-controlled experiments be conducted in order to better understand the reliability of this approach when applied at large spatial scales.

While we developed our model in the context of an eDNA release experiment with a single source, eDNA production at a site can be conditioned on latent occupancy states. This model can also be used for occupancy-type designs with independent sites, with site concentration conditioned on latent occupancy states. A principal implication of our model for natural resource management is that for occupancy analyses, site, sample, and replicate variability in concentration and copy number induce detection heterogeneity at all 3 of these levels. If this variability is not modeled using either observed covariates or random effects, ecological inference may be unreliable. Perhaps the most important effect is that if 1) sites vary in concentration and 2) at least some site concentrations are low enough to lead to missed detections, there will be site heterogeneity in detection probability and subsequent underestimation of the occupancy probability, and thus the potential for misguided natural resource decision making (Royle and Nichols, 2003). Also of importance is that the reliability of inference from false positive occupancy models depends on how well heterogeneity in true positive detection is accounted for (McClintock et al., 2010; Ferguson et al., 2015).

Replacing our eDITH process with processes for independent site occupancy and concentration conditioned on site occupancy would allow for the occupancy states and concentrations at unmeasured sites to be jointly estimated. A caveat, though, is that the reliability of the concentration estimates at sites with no detections (disproportionately those with the lowest concentrations) will likely depend on the adequacy of the distribution used to model site concentration variability (e.g., lognormal variability in site concentration). Further, the performance of such a model will depend on the abundance and reliability of copy number data. Even if not used in practice for occupancy analyses, our model can be used to assess how reliable ecological inference is in the presence of these forms of detection heterogeneity. Ultimately, biostatistical approaches such as those presented in this study help resolve uncertainties associated with eDNA fate and transport as well as qPCR detection and measurement of scarce molecules. Implementation of these types of approaches is needed to more fully take advantage of the potential for eDNA analysis to contribute natural resource management (Kelly et al., 2023).

## Open Research Statement

Data and code are provided at: https://datadryad.org/stash/share/b7xEF9gC-YkK4W5uzf89dF2EwiABZOsCnquuMcbjqE

## Acknowledgments

We thank Diogo Bolster for recommending the power law removal model over the Weibull to provide a similar fit with one less parameter and Andy Royle for reviewing a previous version of this manuscript. Any use of trade, firm, or product names is for descriptive purposes only and does not imply endorsement by the U.S. Government.

## Author Contributions

AS and PH conceived and designed the experiment. BA led model development, data analysis, and the writing of the manuscript. PH and AS contributed to model development and data analysis. PH performed all laboratory analyses. All authors contributed to data collection and writing the manuscript.

## Appendix A MCMC Details

Here, we will describe the priors we used for each model parameter and the MCMC details for parameter updates in the inhibitor models that required different MCMC samplers than the Nimble software can supply.

### Priors

The list of priors for each model can be found in Table A1. Our general strategy was to prefer uninformative priors where feasible, truncate priors to reflect prior knowledge when multimodality is present, truncate priors to improve convergence time, and use moderately informative priors for parameters with binary responses due to sparse data and/or potential separation (e.g., Clipp et al., 2021). The prior knowledge to prevent multimodality we incorporated was 1) we assumed the probability of inhibition must either be unrelated to the number of copies in a replicate or decline as a function of copy number and 2) we assumed the detection probability must either be unrelated to the number of copies in a replicate or increase as a function of copy number. To reduce the time to convergence for the inhibitor model in the simulation study, we truncated the prior for *θ*_0_ such that we only considered that the thinning rate was in the interval (0.03 - 0.82), and we truncated the prior for *σ*^*rep*^ lower than for the null models.

**Table A1.** A list of prior distributions used for each of the models we considered, inhibitor-power law (I-PL), inhibitor-exponential (I-E), null-power law (N-PL), and null-exponential (N-E). “U” indicates “uniform,” “Norm” indicates “Normal,” and “ST” indicates “Student’s T.” “T(a,b)” indicates lower truncation at a and upper truncation at b.

| Par | Model |  |  |  |
| --- | --- | --- | --- | --- |
|  | I-PL | I-E | N-PL | N-E |
| $p_0$ | U(0,50000) | U(0,50000) | U(0,50000) | U(0,50000) |
| $\lambda_{site}$ | U(0,2) | . | U(0,2) | . |
| $\tau$ | . | U(0,5000) | . | U(0,5000) |
| $\theta_{site}$ | U(0,1) | U(0,1) | U(0,1) | U(0,1) |
| $\sigma_{samp}$ | U(0,5) | U(0,5) | U(0,5) | U(0,5) |
| $\beta_0^w$ | Logistic(0,1) | Logistic(0,1) | . | . |
| $\beta_1^w$ | Norm(0,sd=5)T(-∞,0) | Norm(0,sd=5)T(-∞,0) | . | . |
| $\beta_2^w$ | U(0,50000) | U(0,50000) | . | . |
| $\sigma^w$ | ST(0,sd=1,df=4)T(0,∞) | ST(0,sd=1,df=4)T(0,∞) | . | . |
| $\theta_0$ | Logistic(0,1)T(-3.5,1.5) | Logistic(0,1)T(-3.5,1.5) | . | . |
| $\gamma_0$ | Logistic(0,1) | Logistic(0,1) | Logistic(0,1) | Logistic(0,1) |
| $\gamma_1$ | Norm(0,sd=5)T(0,∞) | Norm(0,sd=5)T(0,∞) | Norm(0,sd=5)T(0,∞) | Norm(0,sd=5)T(0,∞) |
| $\sigma^{rep}$ | U(0,0.6) | U(0,0.6) | U(0,1) | U(0,1) |

### MCMC Details

We required two custom MCMC updates for the Inhibitor Models, one for the replicate inhibition states, ***w*** and a second for the sample concentrations, ***C***^*sample*^. Additionally, we added a second update for the replicate inhibition states to further improve mixing. We will describe the updates for the replication inhibition states, followed by the update for sample concentrations.

The default update for *w*_*i,j,k*_ assigned by Nimble is a Binary Sampler that computes and samples from the distribution

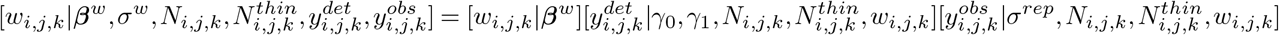

where *w*_*i,j,k*_ sets 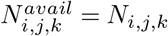 when *w*_*i,j,k*_ = 0 and 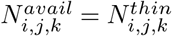 when *w*_*i,j,k*_ = 1. But in order to accept an update from an inhibited state that is consistent with the observed data to an uninhibited state that is consistent with the observed data, and vice versa, you will often need to jointly propose *w*_*i,j,k*_, *N*_*i,j,k*_, and 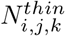 together so that the proposed 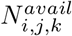 remains consistent with the observed data. This becomes more important as the copy number in replicates increases and the difference between *N*_*i,j,k*_ and 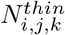 is greater. We used two Metropolis-Hastings samplers to accomplish this in slightly different ways.

#### Update 1

For the first *w*_*i,j,k*_ sampler, we use the following proposal algorithm for a Metropolis-Hastings update.

- Propose 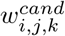 from 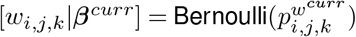
- Propose 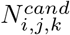 from 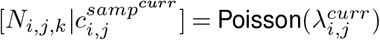, where 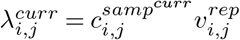
- Propose 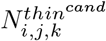 from 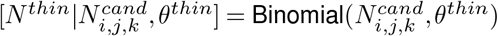
- Set 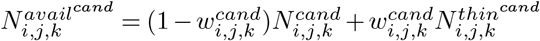
- Update 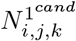 given 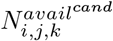
- Update 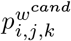 and 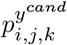 given 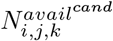

By proposing these candidates from their prior distributions, the proposal probabilities cancel with the likelihoods except for the likelihood of the observed data, leaving us with the Metropolis-Hastings ratio

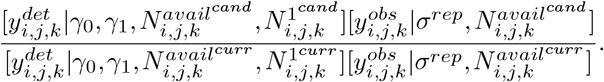

#### Update 2

For the second *w*_*i,j,k*_ update, we use a proposal that does not change the observation model likelihood and may be more likely to be accepted for some latent variable configurations that are unlikely to be accepted in Update 1.

- If 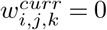
  - Propose 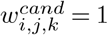
  - Propose 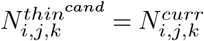 (so the observation model likelihood does not change)
  - Propose 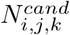 from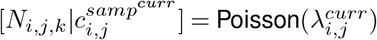, where 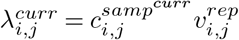
  - Set 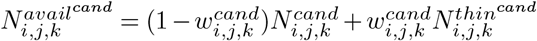
  - Update 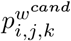 given 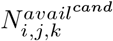

We call this the “shift up” proposal because we propose to raise both the true and thinned copy number. Similarly, we use a “shift down” proposal in the reverse direction.

- If 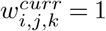 1
  - Propose 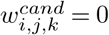
  - Propose 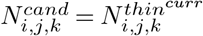(so the observation model likelihood does not change)
  - Propose 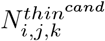 from 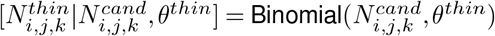
  - Set 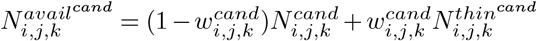
  - Update 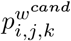 given 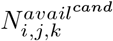

For each case, the Metropolis Ratio is

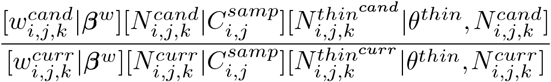

Since the proposals are not symmetric except with respect to the inhibition state, the Hastings Ratio for “shift up” is

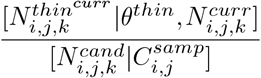

and the Hasting Ratio for “shift down” is

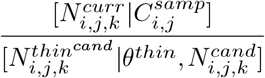

which each cancel different terms in the Metropolis Ratio.

#### Update 3

Updates 1 and 2 are targeted at the individual replication inhibition states. However, the inhibitor model can lead to multimodal posteriors for sample concentrations and we may not be able to move between these modes by updating the replication inhibition states of a sample one at a time. Therefore, we used an update that jointly proposes a new sample concentration, and all latent variables that depend on each sample concentration. We use the following proposal algorithm for each *i* and *j* index on each iteration.

- Propose 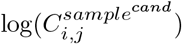 from 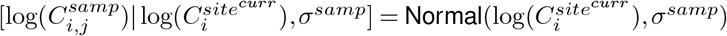
- Update 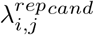 given 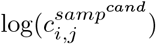
- Propose 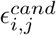 from 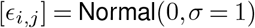
- Propose 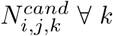 from 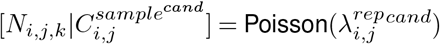
- Update 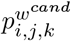 given 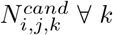
- Propose 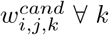 from 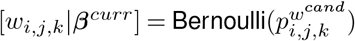
- Propose 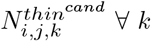 from 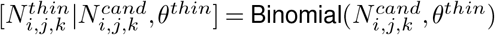
- Set 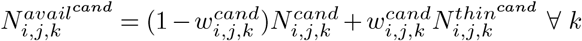
- Update 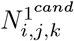 given 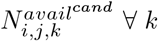
- Update 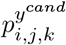 given 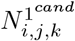 and 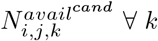

After canceling likelihoods that cancel with proposal from priors, the Metropolis-Hastings Ratio is

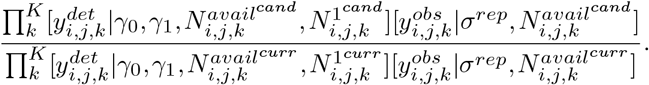

